# Discovery of novel DNA cytosine deaminase activities enables a nondestructive single-enzyme methylation sequencing method for base resolution high-coverage methylome mapping of cell-free and ultra-low input DNA

**DOI:** 10.1101/2023.06.29.547047

**Authors:** Romualdas Vaisvila, Sean R. Johnson, Bo Yan, Nan Dai, Billal M. Bourkia, Minyong Chen, Ivan R. Corrêa, Erbay Yigit, Zhiyi Sun

**Author notes:** These authors contributed equally (Romualdas Vaisvila, Sean R. Johnson).

## Abstract

Cytosine deaminases have important uses in the detection of epigenetic modifications and in genome editing. However, the range of applications of deaminases is limited by a small number of well characterized enzymes. To expand the toolkit of deaminases, we developed an in-vitro approach that bypasses a major hurdle with their severe toxicity in expression hosts. We systematically assayed the activity of 175 putative cytosine deaminases on an unprecedented variety of substrates with epigenetically relevant base modifications. We found enzymes with high activity on double- and single-stranded DNA in various sequence contexts including novel CpG-specific deaminases, as well as enzymes without sequence preference. We also report, for the first time, enzymes that do not deaminate modified cytosines. The remarkable diversity of cytosine deaminases opens new avenues for biotechnological and medical applications. Using a newly discovered non-specific, modification-sensitive double-stranded DNA deaminase, we developed a nondestructive single-enzyme 5-methylctyosine sequencing (SEM-seq) method. SEM-seq enables accurate, high-coverage, base-resolution methylome mapping of scarce biological material including clinically relevant cell-free DNA (cfDNA) and single-cell equivalent 10 pg input DNA. Using SEM-seq, we generated highly reproducible base-resolution 5mC maps, accounting for nearly 80% of CpG islands for a low input human cfDNA sample offering valuable information for identifying potential biomarkers for detection of early-stage cancer and other diseases. This streamlined protocol will enable robust, high-throughput, high-coverage epigenome profiling of challenging samples in research and clinical settings.

## Introduction

Cytosine deaminases are widespread enzymes that are involved in numerous important cellular processes. In eukaryotes, the APOBEC (Apolipoprotein B mRNA Editing Catalytic Polypeptide-like) family of proteins plays important roles in antibody diversification and innate immunity against retroviruses^1^. In bacteria, deaminases are found in polymorphic toxin systems (PTS), which are multi-domain proteins involved in intra- and interspecies competition^2–4^. Deaminases which act on nucleic acids have been used in many biotechnological applications, including as key components of base editing tools^5–7^ and in assays to detect various DNA and RNA modifications^8–12^.

Recently, APOBEC3A deaminase was featured as a non-destructive enzymatic alternative to bisulfite conversion to detect 5-methylcytosine (5mC) and 5-hydroxymethylcytosine (5hmC) DNA modifications (EM-seq^9^ , LR-EM-seq^10^ and ACE-seq^11^). APOBEC3A deaminates cytosine (C) to uridine (U) in single-stranded DNA (ssDNA). APOBEC3A also deaminates 5mC and 5hmC, albeit with reduced activity, but does not deaminate 5-carboxylcytosine (5caC) and glucosylated 5hmC (5gmC). In ABOBEC3A-based modification detection assays, 5mC can be protected through conversion to 5caC and 5gmC by combined Tet methylcytosine dioxygenase 2 (TET2) and T4-phage beta-glucosyl transferase (T4-BGT) activity.

Compared to bisulfite conversion-based sequencing methods^13^, enzymatic deamination protocols do not damage DNA, require lower amounts of input DNA, produce less biased data, and are more compatible with long read sequencing and enrichment of long amplicons^9–11^. We envisioned that the discovery of deaminases with new properties would open the possibility of leaping beyond some of the limitations of current enzymatic methods. Specifically, sequence-agnostic robust activity on double-stranded DNA (dsDNA) combined with a lack of activity on 5mC and 5hmC would enable a streamlined, one-step, one-enzyme protocol for 5mC mapping.

About a decade ago, Iyer, Zhang, and colleagues reported extensive computational analyses of the diversity and phylogenetic distribution of enzymes with the deaminase fold, including a large variety of putative cytosine deaminases found in bacterial polymorphic toxin systems^2,3^. For many years, most of that sequence space remained unexplored experimentally. Recently, Mok, de Moraes, and colleagues described the characterization, engineering, and application to base editing of DddA, of a cytosine deaminase from bacterial toxin systems acting on dsDNA substrates^5–7^. We considered DddA unsuitable for epigenetics modification detection due to its overall low activity and strong sequence preference. Nevertheless, inspired by these earlier works, we endeavored to discover and characterize a broader range of putative cytosine deaminases and assess their suitability for cytosine modification detection and other applications.

In this study, we report an extensive survey of the enzymatic activity of cytosine deaminases from bacterial polymorphic toxin systems, phages, and gene cassettes. We expressed deaminases using an in-vitro system that circumvents their well-documented host toxicity and assayed 175 enzymes from 13 deaminase families. Combining Liquid Chromatography–Mass Spectrometry (LC-MS) and Next Generation Sequencing (NGS) methods, we measured their deaminase activity and substrate selectivity on ssDNA, dsDNA, using unmodified and modified substrates incorporating 5-methylcytosine (5mC), 5-hydroxymethylcytosine (5hmC), 5-formylcytosine (5fC), 5-carboxycytosine (5caC), 5-glucosyloxymethylcytosine (5gmC), and N4-methylcytosine (N4mC). Our work uncovered bacterial deaminases with diverse activities, including enzymes with a wide range of sequence preferences and modification sensitivities on ssDNA and dsDNA. We identified a subset of enzymes with properties desirable for improving epigenetic modification detection methods. Most notably, we found a modification-sensitive deaminase that is active on dsDNA without sequence constraints. We demonstrated the application of this enzyme in a new one-tube deaminase-mediated sequencing method, SEM-seq, for human methylome profiling.

## Results

### In-vitro screening of 175 deaminase sequences

We developed a bioinformatics strategy to look for putative DNA deaminase sequences from public databases. Then, we devised an experimental pipeline to test deamination activity in-vitro (Fig. 1a).

**Figure 1.**
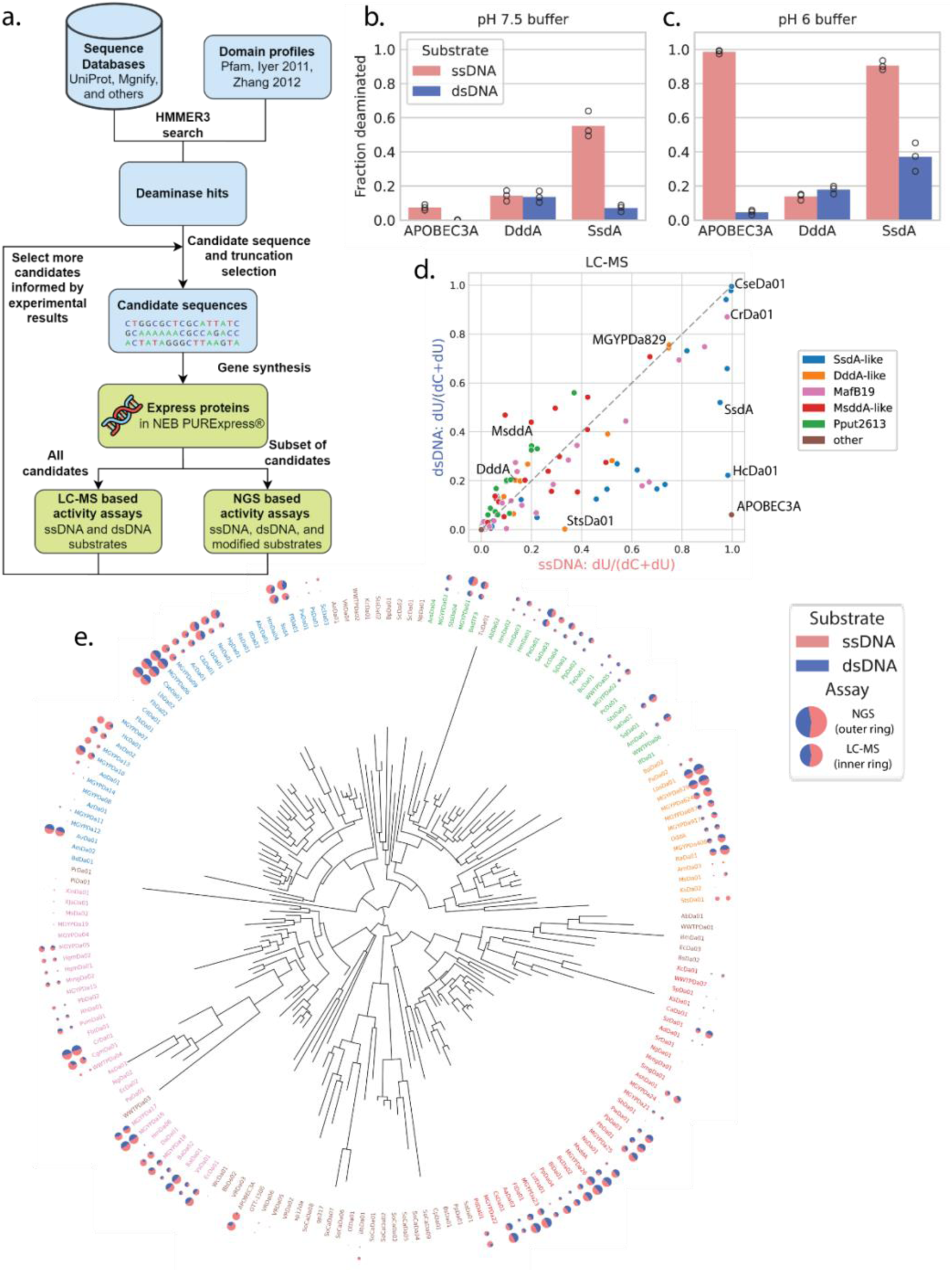
Screen design and results overview. a) Schematic of candidate selection and screening strategy. b) Activity of control enzymes expressed in PURExpress and assayed in pH 7.5 buffer. c) Activity of control enzymes expressed in PURExpress and assayed in pH 6 buffer. d) Activities of all screened enzymes, as measured by the LC-MS assay. e) Maximum likelihood tree of screened enzymes with activities on unmodified ssDNA (pink) and dsDNA (blue), as measured by the LC-MS assay (inner ring) and NGS assay (outer ring) indicated by area on pie charts. Enzymes that showed low activity in the LC-MS assay were not assayed by NGS. DNA emoji designed by OpenMoji – the open-source emoji and icon project. License: CC BY-SA 4.0.

Using as queries, HMMER3^14^ deaminase family profiles from Pfam^15^ and reports by Iyer, Zhang and colleagues^2,3^ (Supplementary Fig. 1), we searched for new deaminases in six different databases, with most candidates coming from UniProt^16^ or Mgnify^17^. We picked hits from diverse deaminase families with the intent to cover a broad range of the sequence space and thus catalog deaminase activities. Candidate selection was performed over many iterations. Active enzymes in our initial screen primarily derived from bacterial polymorphic toxin systems. Later rounds focused on sequences in the same families as those of the active enzymes from previous selections.

Because of their inherent toxicity to expression hosts, deaminases are often obtained in low yields, or are subjected to mutations that inactivate enzymatic activity. As such, putative deaminase proteins were expressed using an in-vitro protein synthesis system. An LC-MS-based assay was developed to quantify deaminase activities on unmodified cytosines in ssDNA, dsDNA, and RNA substrates. The single-stranded ΦX174 Virion DNA and the double-stranded ΦX174 RF I DNA, sharing the same template sequence were used as DNA substrates. These 5,386-nucleotide (nt) length substrates present C in a rich diversity of sequence contexts, thus allowing the detection of deaminase activities regardless of the sequence specificities. Isotope labeled firefly luciferase (Fluc) mRNA was used as the substrate for testing deaminase activity on RNA for a subset of deaminases.

We validate our screening assay on the previously published SsdA and DddA enzymes. Considering the potential mutagenicity of DNA deaminases, we compared the use of the PURExpress in-vitro protein synthesis system with different transcription templates. These were composed of either unmodified DNA or of DNA with 5hmdC in place of dC to potentially block template mutation during the transcription and translation reactions. Of note, DNA templates constructed from the 5hmdCTP and dCTP generated similar results (Supplementary Fig. 2a). Further investigation showed that deaminase activity is inhibited by the 1X PURExpress buffer (Supplementary Fig. 2b) but can be rescued in the diluted buffer. Our data suggested this is a viable approach to efficiently produce active proteins of toxic deaminases for further characterization. As a case in point, we showed by LC-MS analysis that SsdA, like APOBEC3A, prefers ssDNA, and DddA is active on dsDNA (Fig. 1b and Supplementary Fig. 2 c,d), which is in agreement with the published results. We further uncovered that all these three enzymes have equal or better activity in a lower pH buffer (pH 6)^18^ than that of a previously published buffer (pH 7.5)^7^ (Fig. 1b,c).

We screened a total of 175 new enzymes across 13 cytosine deaminase families (Fig. 1d,e, Supplementary Fig. 1). Our assay revealed that many bacterial DNA deaminases act on both ssDNA and dsDNA, with some deaminating both types of substrates with equal efficiency. Five deaminase families, DddA-like, SsdA-like, Pput2613, MafB19, and MsddA-like, accounted for nearly all enzymes with DNA activity detectable by LC-MS (Fig. 1d,e). In four of the five families, at least one deaminase was found with higher RNA deaminase activity than APOBEC3A (Supplementary Fig. 3), which has previously been reported to have activity on RNA substrates^19^. Examples of enzymes highly active on DNA were CseDa01, MGYPDa06, and LbDa02, which deaminated close to 100% of cytosines in dsDNA and ssDNA substrates;

CrDa01, which deaminated 87% and 97% of cytosines in dsDNA and ssDNA, respectively; and MGYPDa829 and LbsDa01, which deaminated about 75% of cytosines in both dsDNA and ssDNA substrates. When we tested at lower concentrations, MGYPDa829 (DddA-like) showed equal activity on ssDNA and dsDNA, whereas CseDa01 (SsdA-like) preferred ssDNA (Supplementary Fig. 4). Overall, most of the active bacterial deaminases display a preference for ssDNA substrates. While we found a few cytosine deaminases that strongly prefer single-stranded substrates, such as HcDa01 and StsDa01, we did not find a single bacterial deaminase that only acts on dsDNA. For simplicity, herein we will use the term “dsDNA deaminase” for enzymes that act on dsDNA, even though they also deaminate ssDNA to some extent.

### A wide spectrum of sensitivity to DNA cytosine modifications

We next measured deaminase activity on a variety of physiologically relevant cytosine modifications. We developed an NGS assay to characterize deaminase sequence preference and sensitivity to 5C-methylation, 5C-hydroxymethylation, 5C-glucosylation, and N4-methylation (N4mC) in both single-stranded and double-stranded DNA. In addition, we used the LC-MS assay to examine activity on 5C-formylated, and 5C-carboxylated substrates. To validate these methods, we showed that the deamination efficiency on unmodified substrates measured by NGS was broadly consistent with that of LC-MS (Pearson r = 0.85 and 0.86 for ssDNA and dsDNA respectively) (Supplementary Fig. 5).

We found that the screened enzymes generally had low activity on N4mC (Supplementary Fig. 6) but displayed a wide spectrum of activities on different modification types at the cytosine 5-position (Fig. 2, Supplementary Fig. 7). We clustered the deaminases into five functional clusters based on their specificity for C, 5mC, 5hmC and 5gmC in dsDNA. Most deaminases showed a moderate decrease of activity toward 5mC and 5hmC modifications, and a strong decrease of activity toward 5gmC compared to unmodified dsDNA substrates (Fig. 2a,b, clusters #2 and 4). A subset of deaminases displayed similar activities across all the cytosine types (Fig. 2a,b, cluster #1). This group includes CseDa01, which efficiently deaminates C, 5mC, 5hmC, and even gmC to nearly 100%. Another interesting subset showed decreased activity on 5mC and 5gmC, but not on 5hmC (Fig. 2a,b, cluster #3). This subset, which includes PfDa01 and SsdA, can potentially be used to distinguish 5hmC from 5mC. Enzymes from clusters #1-4 tended to have more relaxed modification sensitivity to ssDNA compared to dsDNA (Supplementary Fig. 7). The most exciting discovery was of a unique functional cluster with very low or no activity on modified substrates (Fig. 2a,b, cluster #5). The enzymes from cluster #5 with the highest deamination activity of unmodified C were found to be phylogenetically related (Fig. 1e). We referred to the most active enzyme of this group (92% of unmodified cytosine deaminated and less than 3% of the modified cytosines), originating from human mouth metagenome (European Nucleotide Archive accession ERZ773077)^20^, as “Modification-Sensitive DNA Deaminase A” (MsddA) and the family of deaminases related to it as MsddA-like.

**Figure 2.**
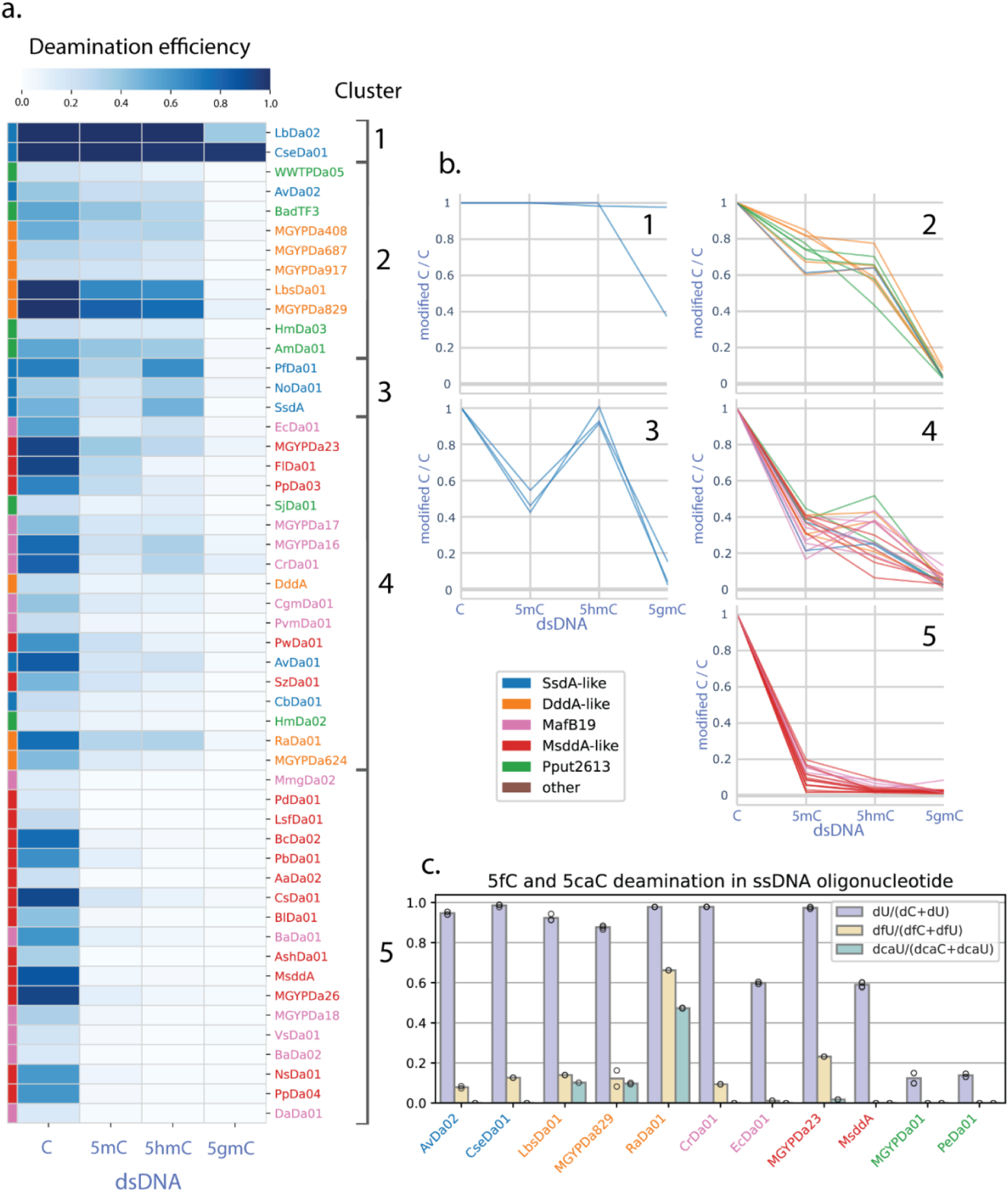
Deamination efficiency of representative enzymes on C-5 modified substrates in dsDNA. a) Deamination efficiency on C, 5mC, 5hmC, and 5gmC in dsDNA, measured by NGS assay. The enzymes were grouped into five clusters, using average linkage clustering of cosine distances between deamination efficiencies. b) Deamination efficiency on dsDNA C, 5mC, 5hmC, and 5gmC divided by deamination efficiency on C of enzymes in the five clusters. c) Deamination efficiency on 5fC and 5caC in ssDNA oligonucleotides by representative deaminases from each family, measured by LC-MS assay.

We also examined deamination activities on 5fC and 5caC of representative enzymes from each family using ssDNA oligonucleotide substrates containing four cytosines in a ‘Cg’ context in which all cytosines were either unmodified, 5-formylated, or 5-carboxylated (Fig. 2c, Supplementary Fig. 8). We observed that MsddA, EcDa01 (MafB19 family), and the two enzymes from the Pput2613 family did not deaminate either modified substrate. In contrast, all other tested deaminases displayed some activity on 5fC, and the three enzymes from the DddA-like family had activity on both 5fC and 5caC.

### A wide variety of DNA sequence preferences

The single-base resolution of our NGS assay and the use of a large size and sequence complexity of substrates enabled us to accurately survey the sequence preference landscape for our deaminase library. Sequence Logo plotting^22^ of 10-base context on either side of sites deaminated in unmodified dsDNA uncovered a diversity of context preferences (Fig. 3a-c, Supplementary Fig. 9), including a subset of enzymes that display no sequence constraints (Fig. 3a). The non-specific enzymes also tend to have high overall activity (Fig. 3d). In addition to displaying different preferences for sequences at the 5’ end position(s) to the deaminated cytosine, reminiscent of enzymes from the eukaryotic APOBEC deaminase family^18^, many bacterial deaminases display sequence selectivity for the immediate 3’ position, some specifically recognizing CpG dinucleotides (Fig. 3b,d). Deaminase recognition sequence sites can extend beyond the nCn context, with preferences for sequences of various lengths and compositions (Fig. 3c).

**Figure 3.**
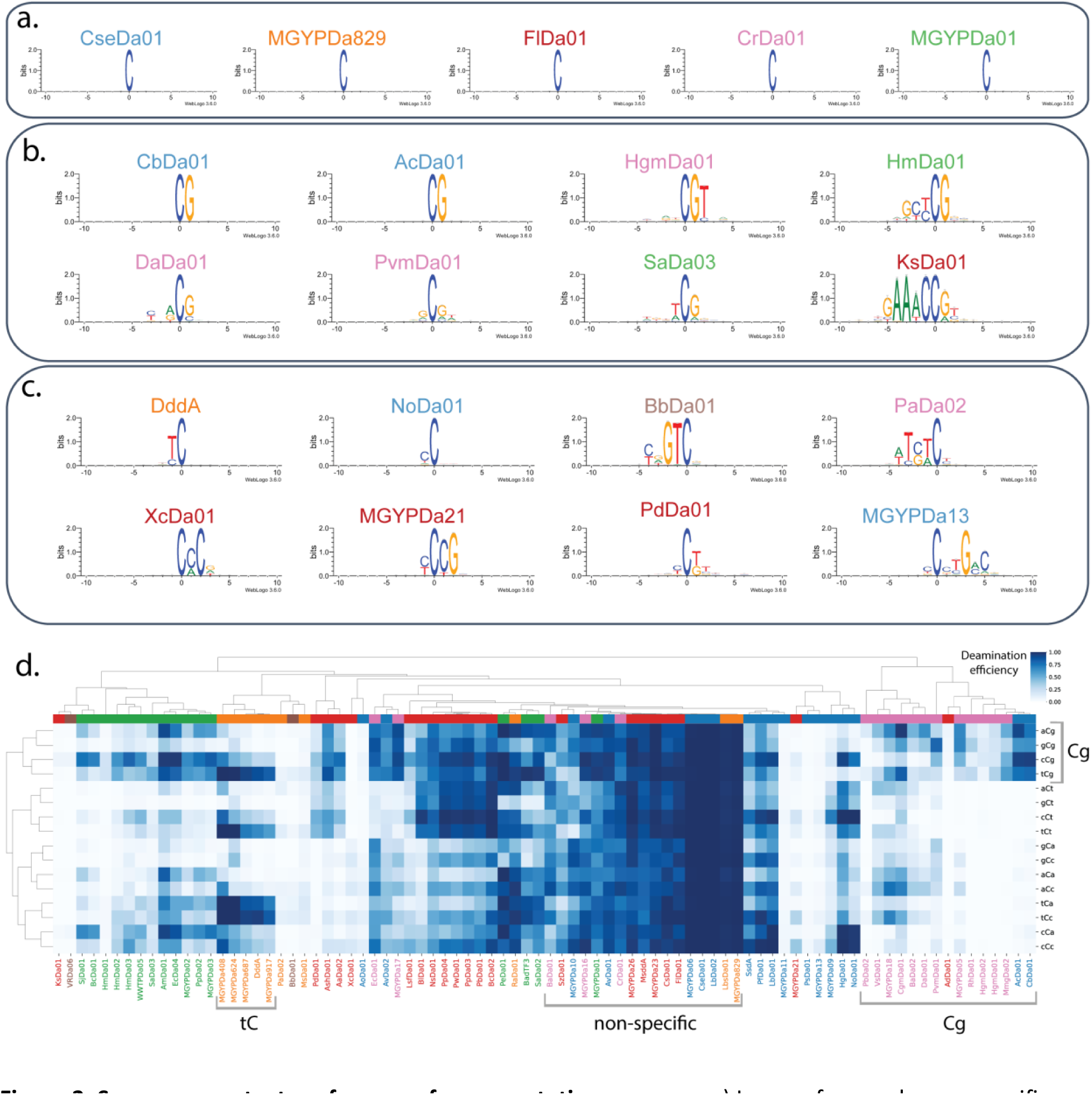
Sequence context preference of representative enzymes. a) Logos of example non-specific deaminases from each family. b) Logos of example deaminases with preference for G at the +1 position, including CpG-specific deaminases. c) Logos of example deaminases with diverse sequence specificities. d) Activity on representative deaminases in the nCn contexts of unmodified dsDNA. Rows and columns are sorted based on average linkage clustering of cosine distances. All logos are of sites with >=50% deamination efficiency in unmodified dsDNA. Deamination efficiencies measured by NGS assay. Names are colored according to family: SsdA-like (blue), DddA-like (orange), MafB19 (pink), MsddA-like (red), Pput2613 (green), other (brown).

As a more systematic analysis of sequence preference across different substrate types, we calculated the average deamination efficiency of all nCn sequences in unmodified and modified dsDNA and ssDNA substrates and conducted clustering analysis across diverse enzymes and specificities (Fig. 3d, Supplementary Fig. 10). We observed that enzymes from the same family tend to cluster together. For example, MafB19 deaminases tend to prefer Cg, DddA-like deaminases tend to prefer tC, and MsddA-like deaminases tend to have reduced activity on Ca. However, all five families contained examples of non-specific deaminases (Fig. 3a). Specificities were generally consistent across substrates, noting that enzymes tend to display more context specific preferences on substrates on which they had lower total activity (Supplementary Fig. 10).

### SEM-seq, a nondestructive streamlined single-enzyme methylation sequencing method for accurate base-resolution methylome analysis

The discovery of the non-specific methylation-sensitive dsDNA deaminase MsddA enabled us to perform SEM-seq, a single-enzyme method for methylation sequencing at single base resolution. SEM-seq eliminated the need for TET2/T4-BGT protection and denaturing steps that are required of APOEC3A-based protocols (Fig. 4a).

**Figure 4.**
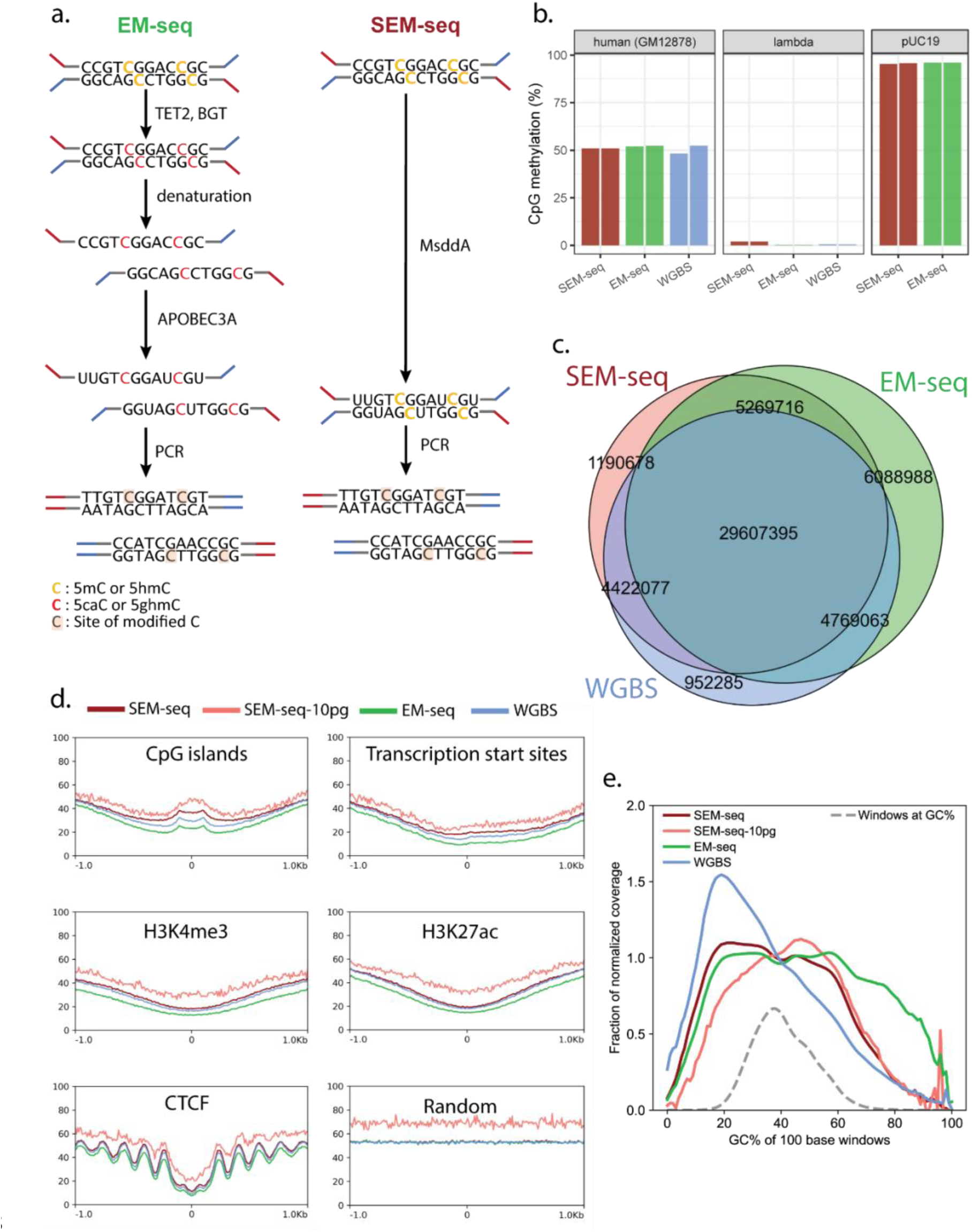
SEM-seq Human GM12878 DNA compared to EM-seq, and whole genome bisulfite sequencing (WGBS). a) Schematic of EM-seq and SEM-seq protocols. b) Total CpG methylation in GM12878, lambda (negative internal control) and CpG-methylated pUC19 (positive internal control) DNA. c) Counts of methylated CpG sites called in common by the three methods. d) Comparison of CpG methylation profiles in neighborhoods around specific genomic features. e) Effects of GC-content on read coverage from the three methods.

We validated SEM-seq using genomic DNA from the human GM12878 cell line and benchmarked it against two published datasets from standard EM-seq^9^ and WGBS (ENCODE accession ENCSR890UQO)^23^ protocols. Unmethylated lambda and CpG-methylated pUC19 DNA were spiked in as controls to estimate deamination rates. Two technical replicates of SEM-seq were sequenced on the Illumina Nova-seq platform. 277 million paired-end reads from two replicates of each method were used for the comparative analyses.

The library quality metrics and methylation results were highly consistent between technical replicates, suggesting a high reproducibility and robustness of SEM-seq (Supplementary Fig. 11). SEM-seq libraries produced very similar CpG methylation results for human GM12878 as those of EM-seq and WGBS (Fig. 4b-d), despite a slightly lower CpG deamination rate on unmodified cytosines for lambda phage DNA (SEM-seq 98.0%, EM-seq 99.8%; WGBS 99.4%). The 5mC non-conversion rates calculated from the CpG-methylated pUC19 DNA were 95.4% and 95.7% for each of the two SEM-seq replicates. These data were nearly identical to that of EM-seq (∼96%) (Fig. 4b), but without requiring protection by TET2 and BGT. The conversion rates of unmodified pUC19 CH sites were 96.3% and 95.7% for the two SEM-seq replicates. Due to generally low methylation level of CH sites in the human genomes, we focused our methylome analysis on CpG sites. After combining the data of the two technical replicates, SEM-seq identified 40.4 million high-confidence methylated CpG sites, of which 86% and 84% agreed with EM-seq and WGBS datasets, respectively (Fig. 4c). The methylation levels of commonly methylated CpG sites are also highly correlated between SEM-seq and the other two methods (Pearson correlations: SEM-seq vs. EM-seq 0.91, SEM-seq vs. WGBS 0.88, EM-seq vs. WGBS 0.90). SEM-seq accurately produced the expected CpG methylation profiles for key epigenomic features, such as an enrichment of methylation in CpG islands, a reduced methylation levels near transcription start sites and at active chromatin markers and enhancers, and a typical regularly spaced oscillation pattern surrounding CTCF binding sites with depleted methylation at the center^24^ (Fig. 4d). Furthermore, SEM-seq, like EM-seq, is non-destructive and gives more even read coverage across genomic regions of variable GC contents than the bisulfite-based WGBS method (Fig. 4e).

### SEM-seq for cfDNA and 10 pg input

The streamlined, robust protocol of SEM-seq makes it an advantageous method for obtaining accurate and highly reproducible methylome information from scarce biological samples such as cell free DNA (cfDNA), which has important clinical applications for noninvasive prenatal diagnosis and early cancer detection and monitoring^25^. We applied SEM-seq to human cell free DNA (cfDNA) and sequenced two replicate SEM-seq libraries each using 3 ng of cfDNA, generating 187.0 million and 204.5 million paired-end reads. The two cfDNA libraries covered 26.6 million and 30.6 million CpG sites respectively with at least 3X coverage. Among them, 25.6 and 29.4 million methylated CpG sites were identified with 18.5 million agreeing between the two replicates (Supplementary Fig. 12a). The methylation levels of individual CpG sites also correlated well between the replicates (Pearson correlation=0.71) demonstrating accurate, base resolution quantification. A single cfDNA SEM-seq library generated accurate 5mC profiles in epigenetic important genomic features resembling those of the human GM12878 genomic DNA (Fig. 5a, Supplementary Fig. 12b). Remarkably, SEM-seq was able to provide single-base-resolution 5mC quantification of 21536 CpG islands accounting for 78% of annotated CpG islands in the human genome, leading to the detection of 11426 hypomethylated (methylation level <20%) and 6712 hypermethylated (methylation level >70%) CpG islands and their associated genes (Supplementary Fig. 12c, Supplementary Table S1). As expected, gene promoter regions are depleted in 5mC globally. SEM-seq allowed us to further examine the distribution and methylation level of individual cytosine sites of each gene at a finer resolution. Consistent with the global trend, many promoter regions are hypomethylated, for example the N-glycanase 1 (NGLY1) promoter (Fig. 5b). However, a subset of genes are heavily modified in the promoter, such as BCL6 corepressor like 1 (BCORL1) (Fig. 5c)—a transcriptional corepressor that was suggested a prognostic factor that promotes tumor metastasis^26^, and LDL receptor related protein 5 like (LRP5L), of which a CpG island near the TSS is also hypermethylated (Fig. 5d). Such high-resolution methylation information is very valuable for detecting aberrant DNA methylation in cancer and other diseased samples and identify potential biomarkers for detection and classification of early-stage cancer.

**Figure 5.**
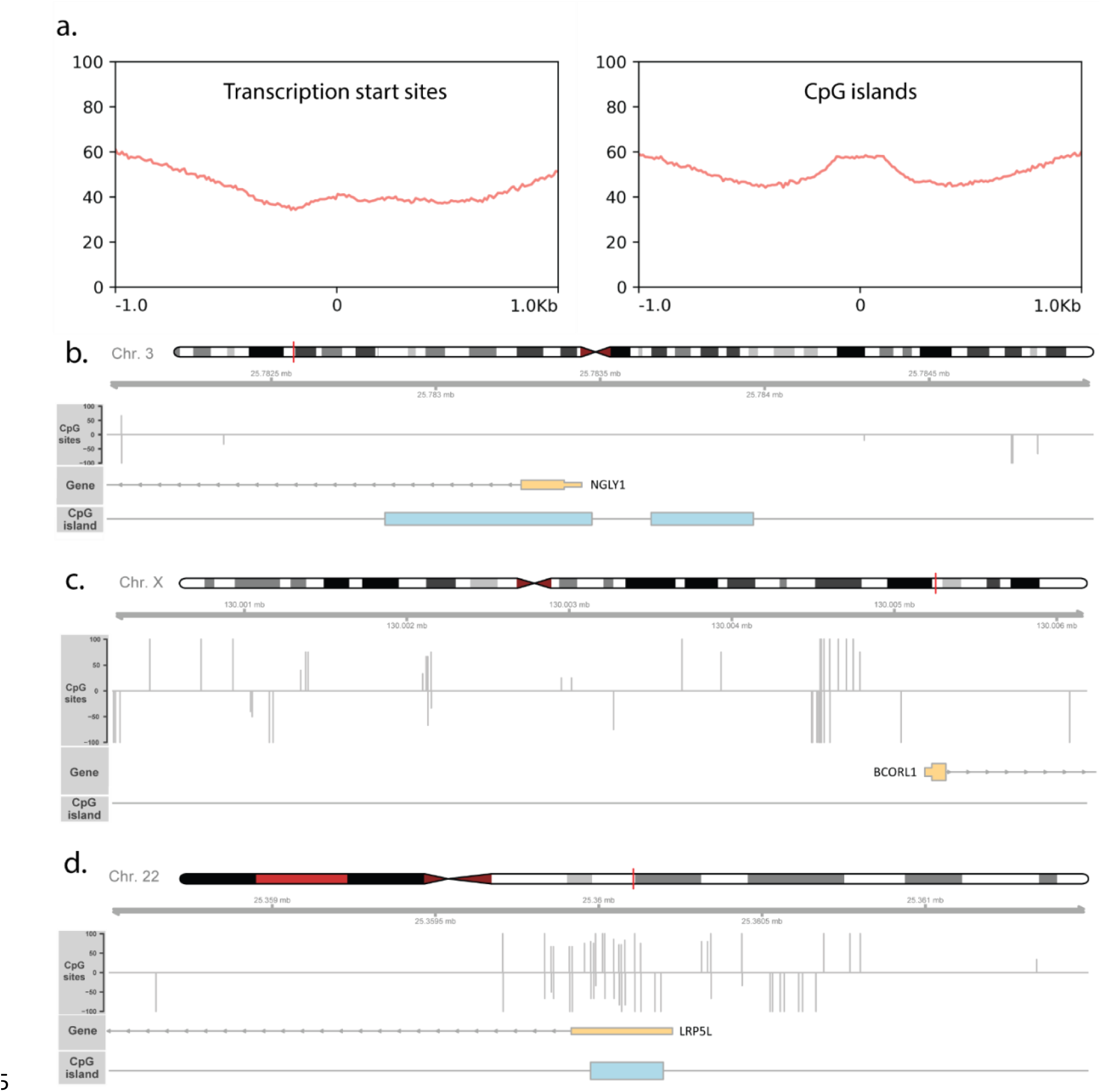
SEM-seq on human cfDNA sample. a) CpG methylation profiles in neighborhoods around transcription start sites (TSS) and CpG islands. b) 1.5 Kb upstream and 1 Kb downstream flanking region of the TSS of NGLY1 (hypomethylated). c) 5 Kb upstream and 1Kb downstream flanking region of the TSS of BCORL1 (hypomethylated). d) 1.5 Kb upstream and 1 Kb downstream flanking region of the TSS of LRP5L (hypomethylated). For b), c), and d), the methylation levels of CpG sites are shown as vertical grey lines. Positive and negative values indicate CpG sites on the top strand and bottom strand, respectively. Coordinates correspond to positions on respective chromosomes (GRch38).

We also made SEM-seq libraries from only 10 pg of human GM12878 DNA, equivalent to single cell input^27^. With 28 million paired-end reads, a single 10 pg library covered 9.3 M CpG sites and 8219 (30%) CpG islands. It revealed similar methylation patterns to the 50 ng EM-seq and SEM-seq libraries, with low GC coverage bias (Fig. 4e), and the expected 5mC distributions at various key genomic features (Fig. 4d). This result suggests that SEM-seq is suitable for single-cell methylome studies.

## Discussion

This work presents a systematic experimental screening of 175 new cytosine deaminases spanning 13 distinct deaminase families. Screened enzymes cover most deaminase families that have been hypothesized to act on cytosines^2,3^. Nearly all the active bacterial deaminases we characterized are found in polymorphic toxin systems (PTS) and come from five of the 13 families: SsdA-like, DddA-like, MafB19, Pput2613, and MsddA-like. The last of which we christened in reflection of the sensitivity to modified cytosines displayed by many of its members.

Our screen reveals that bacterial deaminases are a versatile group of enzymes with diverse and previously unknown properties. We show that many bacterial deaminases act on dsDNA, contrasting with earlier observations that most DNA cytosine deaminases strongly prefer ssDNA substrates. While some bacterial DNA deaminases are strictly single-strand specific enzymes, which parallels all the known eukaryotic DNA deaminases, we did not find any deaminases that only accept double-strand DNA substrates. This suggests that the deamination activity on single-stranded substrates may be the ancestral state of all the bacterial deaminases.

Several recent studies have reported a functional characterization of enzymes from the DddA-like family^28–30^. One study^28^ showed that deletion of SPKK-related motifs at the C-terminus of DddA abrogates its activity on dsDNA. In agreement with those results, we found that StsDa01, a deaminase from the DddA-like family, lacks a C-terminal SPKK-related motif and strongly prefers ssDNA substrates. Furthermore, we found that by truncating the C-terminus of MGYPDa829, this enzyme was converted from having similar activity on both dsDNA and ssDNA to strongly preferring ssDNA (Supplementary Fig. 13). The enzyme resulting from a swap of the C-terminus from MGYPDa829 onto StsDa01 retained a strong preference for ssDNA. We observed that dsDNA deaminases outside the DddA-like family lack SPKK-related motifs, therefore the association between dsDNA activity and the SPKK-related motif does not seem to generalize beyond the DddA-like family.

Bacterial deaminase activities also vary widely across DNA modification types including 5mC, 5hmC, 5fC, 5caC, 5gmC, and N4mC. In contrast to a wide spectrum of activities on modifications at C-5 (Fig. 2), including the first report of deaminases with strong activity on 5caC and 5gmC, the screened enzymes generally had low activity on N4mC (Supplementary Fig. 6). Finally, we found that bacterial deaminases display a broad spectrum of sequence specificities, including enzymes with no apparent biases, enzymes with diverse sequence preferences at the 5’ end position relative to the target C, and enzymes with strong sequence preferences at the 3’ end positions, particularly a strong bias toward CpG dinucleotides.

The diverse properties of bacterial deaminases lend themselves to various applications, including as base editors and for epigenetic modification mapping. Information on substrate preference serves as a key for selecting suitable enzymes for different applications. In base editing applications, their small size (100-200 amino acids) (Supplementary Fig. 14) could provide advantages for therapeutic delivery. Also, a strong ssDNA preference coupled with different context specificities will likely help reduce off-target editing. For detecting the important epigenetic mark 5mC, a sequence independent cytosine deaminase that can discriminate between methylated and unmethylated sites would be ideally suited for broad methylome analysis. Indeed, we demonstrate the successful application in human whole methylome mapping of the non-specific, modification-sensitive dsDNA deaminase MsddA. Our streamlined methylation sequencing protocol, SEM-seq, not only is free of the damage-inducing bisulfite treatment, but also does not require any accessory protection proteins (TET2 and T4-BGT) nor harsh denaturing steps. We leveraged the advantages of SEM-seq by successfully applying it to methylome analysis in clinically relevant cfDNA and in challenging, single-cell equivalent, 10 pg input DNA samples.

Our study unravels another example of the great potential for developing biotechnological tools from previously unexploited bacterial and bacteriophage enzymes, in our case DNA methylation mapping using bacterial DNA deaminase enzymes with diverse substrate preferences and modification sensitivities.

### Limitations of the Study

Deaminases screened in this study or used for the SEM-seq experiments were all produced using the PURExpress in-vitro transcription/translation (IVTT) system without further purification. The lack of purified and quantified deaminases limited our ability to optimize SEM-seq conditions, for example by fine-tuning enzyme concentration. The current SEM-seq conversion rate of unmodified CH is around 96%, which is lower than unmodified CH conversion rates typically observed in EM-seq and WGBS experiments. This CH conversion rate needs to be improved in order to accurately detect low level non-CpG methylations in most mammalian samples. Future work with purified deaminases will focus on optimizing SEM-seq conditions for improved sensitivity of 5mC detection in both CpG and non-CpG contexts.

## Methods

### Candidate selection

To guide the search for new enzymes, we first curated a list of HMMER3^14^ cytosine deaminase sequence profiles. 29 profiles came from the CDA clan (CL0109) from the Pfam^15^ database (version 34) (excluding the TM1506, LpxI_C, FdhD-NarQ, and AICARFT_IMPCHas, which are thought to not encode deaminases), 17 profiles were built from multiple sequence alignments (MSAs) of deaminase families defined by Iyer et al. (2011)^2^, and one profile was built from a multiple sequence alignment found in Zhang et al. (2012)^3^. The profiles from Iyer largely overlap with the Pfam profiles, with a few differences, despite some similar names (Supplementary Fig. 1). The Pfam MafB19-deam (PF14437) profile and the similarly named profile from Iyer were found to be different from each other. None of the enzymes we screened had a best profile match to the Pfam MafB19-deam profile, so our usage of MafB19 corresponds to the Iyer profile.

The Pfam DYW_deaminase (PF14432) profile is biased towards eukaryotic members of that family, whereas the similarly named profile from Iyer captures both eukaryotic and bacterial DYW deaminases. We therefore split the Iyer DYW profile into three separate profiles, the one comprising mostly bacterial enzymes we called SsdA-like, as it contains the previously described SsdA deaminase^7^, the other, which more closely resembles the DYW profile from Pfam, we called Iyer2011_DYW, the combined profile we called Iyer2011_DYW_combined (giving a new total of 18 profiles from Iyer). We found experimentally that the enzymes with strong matches towards the profile from Zhang et al. 2012^3^ tended to have poor deamination activity on modified cytosines. We therefore renamed this profile MsddA-like, for “modification sensitive DNA deaminase A-like”.

Some candidate sequences were selected directly from the MSAs listed in Iyer et al. (2011) and Zhang et al. (2012). Others were selected from hmmsearch hits of the profiles described above against six different databases: UniProt^16^, Mgnify^17^, IMG/VR^31^, IMG/M^32^, gene cassette metagenomes^33^, wastewater treatment plant metagenomes^34^, and GenBank^35^. Candidate selection and experimental screening were performed over many iterations. Candidates were selected from the hmmsearch hits with the intent to cover a broad range of sequence space, but with a focus on sequences similar to those that were shown to be active in earlier rounds. Mok et al. (2020)^5^ and de Moraes et al. (2021)^7^ previously reported on the active bacterial DNA deaminases, SsdA, DddA, and BadTF3, so similarity to those enzymes was also used as a criterion for candidate selection.

Most of the deaminases we tested were found as fusions to larger proteins, for example as parts of polymorphic toxin systems. In our assays we only expressed the deaminase domain rather than the full proteins. To determine the boundaries of the deaminase domain, we ran AlphaFold2^36^ via the LocalColabFold^37^ package on the deaminase domain plus up to about 1000 amino acids upstream and downstream. We visualized the AlphaFold2 predicted structures in PyMOL. N-terminal truncation sites were generally selected at several amino acids before helix 1 of the deaminase domain. The deaminase domains were typically found at the C-terminus of the fusion protein, so for most sequences C-terminal truncations were not necessary. For cases where the boundaries of the deaminase domain were difficult to distinguish, we either tried several different truncations (Supplementary Fig. 15) or relied on the Predicted Aligned Error (PAE) metric reported by AlphaFold2. We found that residues with a low PAE to residues in the core of the deaminase domain corresponded to our intuitive notions of the boundaries of the deaminase domain. To aid in the visualization of PAE, we developed a Colab notebook called PAEView (https://github.com/seanrjohnson/proteinotes/blob/master/colab_notebooks/PAEView.ipynb).

### Sequence naming

For convenience, each screened sequence was given a short name. The names are related to the database or species of origin for the sequence. Da = deaminase, MGYP = Mgnify protein, Hm = hot metagenome, VR = IMG/VR, WWTP = wastewater treatment plant, SoCa = soil gene cassette, chimera = chimeric sequence. Other prefixes are mostly two or three letters drawn from the name of the source organism or the source environment of the metagenome data. Some sequences also have prefixes or suffixes of the form extN#, extC#, d#, Cd#, which indicate, respectively, N-terminal extensions, C-terminal extensions, N-terminal deletions, and C-terminal deletions of the indicated number of residues, compared to the candidate with the un-affixed name.

### Phylogenetic Tree

Amino acid sequence alignments were all calculated using MUSCLE^38^. Trees were generated using raxml-ng (v. 1.1)^39^. The tree was rooted at the midpoint and rendered using ETE3^40^.

### Hmm profile comparison

Similarity between deaminase hmm profiles was computed using hmmer_compare.py, a new python implementation of the algorithm described by Söding^41^. Scores were calculated for each pair of profiles. Scores were converted to distances using the formula:

distance = 1 - (pairwise_score / min(profile1_self_score, profile2_self_score))

A UPGMA tree was generated from the distance matrix using scikit-learn^42^ and rendered with Geneious Prime (https://www.geneious.com/) to visualize distances between profiles.

Code for generating hmmer3 profile trees and alignments is available at: https://github.com/seanrjohnson/hmmer_compare

### Sources of DNA and RNA substrates

Unmodified DNA oligonucleotides were purchased from IDT Coralville, IA), 5caC and 5fC containing oligonucleotides were synthesized by NEB (Ipswich, MA). dsDNA oligonucleotides were annealed in 10 mM Tris–HCl, pH 8.0 buffer*. E. coli* C2566 genomic DNA was purified using Monarch Genomic DNA Purification Kit (NEB Ipswich, MA), GM12878 genomic DNA was obtained from Coriell Cell Repositories (Camden, NJ). Single donor human plasma was obtained from Innovative Research (Novi, MI) and cfDNA was extracted using the BioChain (Newark, CA) cfPure MAX V2 Cell-Free DNA Extraction Kit.

Unmethylated cl857 Sam7 Lambda DNA (Promega, Fitchburg, WI), fully C-methylated XP12 phage DNA (Dr. Yan-Jiun Lee, NEB Ipswich MA), fully C-hydroxymethylated T4147^43^ and fully C-hydroxy-methyl-beta-glucosylated T4 alpha-glucosyl-transferase knockout (AGT-) genomic DNAs (Dr. Lidija Truncaite, Vilnius University Life Sciences Center, Vilnius Lithuania), pRSSM1.PleII plasmid (containing N4mC at cacCgc sites) (Dr. Iain Murray, NEB Ipswich MA) were used as spike in controls in NGS deamination assay.

Isotope labeled firefly luciferase (Fluc) mRNA was synthesized by HiScribe T7 High Yield RNA Synthesis Kit (E2040S, NEB, Ipswich) using stable heavy isotopes of ATP and CTP (NLM-3987-CA and CNLM-4267-CA-20, Cambridge Isotope Laboratory, Tewksbury, MA). Resulting Fluc mRNA was purified twice by Monarch RNA cleanup kit (T2040S, NEB, Ipwsich, MA) to ensure that unincorporated nucleotides were completely removed. RNA was eluted in nuclease-free water, quantified by Qubit RNA broad range assay (Q10211, ThermoFisher, Waltham, MA) and stored at -20 °C.

### In-vitro expression of deaminases

The candidate DNA deaminase genes first were codon-optimized then added flanking sequences containing T7 promoter at 5’ end (GCGAATTAATACGACTCACTATAGGGCTTAAGTATAAGGAGGAAAAAAT) and T7 terminator at 3’ end (CTAGCATAACCCCTCTCTAAACGGAGGGGTTTATTTGG) and ordered as linear gBlocks from IDT (Coralville, IA, USA). Template DNA for in vitro protein synthesis was generated with Phusion® Hot Start Flex DNA Polymerase (NEB. Ipswich MA) using gBlocks as template and flanking primers (Supplementary Table S2). The PCR products were purified using Monarch PCR and DNA Cleanup kit (NEB, Ipswich, MA). DNA concentration was quantified using a NanoDrop spectrophotometer (Thermo Fisher Scientific, Inc., Waltham, MA). 100 - 400 ng of PCR fragments were used as template DNA to synthesize analytic amounts of DNA deaminases using PURExpress In Vitro Protein Synthesis kit (NEB, Ipswich, MA) following manufacturer’s recommendations.

### Deamination activity assay on single and double stranded ΦX174 DNA substrates

To test the activity of in-vitro expressed DNA deaminases, 2 μL of PURExpress sample was mixed with 300 ng of ΦX174 Virion DNA (ssDNA substrate, NEB Ipswich MA) or ΦX174 RF I DNA (dsDNA substrate, NEB, Ipswich MA) into buffer containing 50 mM Bis-Tris pH 6.0, 0.1% Triton X-100 and incubated for 1 h at 37°C for a total volume of 50 μL. The deaminated ΦX174 DNA was purified using Monarch PCR and DNA Cleanup kit (NEB, Ipswich, MA). DNA concentration was quantified using a NanoDrop spectrophotometer (Thermo Fisher Scientific, Inc., Waltham, MA). 150 ng of deaminated DNAs were digested to nucleosides with the Nucleoside Digestion Mix (NEB, Ipswich, MA) following manufacturer’s recommendations. LC-MS/MS analysis was performed by injecting digested DNAs on an Agilent 1290 Infinity II UHPLC equipped with a G7117A diode array detector and a 6495C triple quadrupole mass detector operating in the positive electrospray ionization mode (+ESI). UHPLC was carried out on a Waters XSelect HSS T3 XP column (2.1 × 100 mm, 2.5 µm) with a gradient mobile phase consisting of methanol and 10 mM aqueous ammonium acetate (pH 4.5). MS data acquisition was performed in the dynamic multiple reaction monitoring (DMRM) mode. Each nucleoside was identified in the extracted chromatogram associated with its specific MS/MS transition: dC [M+H]^+^ at m/z 228.1→112.1; dU [M+H]^+^ at m/z 229.1→113.1; d^m^C [M+H]^+^ at m/z 242.1→126.1; and dT [M+H]^+^ at m/z 243.1→127.1. External calibration curves with known amounts of the nucleosides were used to calculate their ratios within the samples analyzed.

### Deamination activity assay on single and double-stranded oligonucleotide DNA substrates

To test the activity of in vitro expressed DddA and SsdA DNA deaminases on oligonucleotide substrates (Supplementary Table S3), a 2 µL of PURExpress sample was mixed with 300 ng of 44 bp single-stranded or double-stranded DNA substrates (annealed in 10 mM TRIS ph=8.0) with single cytosine in the contexts TC, AC, CC, and GC in a buffer consisting of 20 mM Tris-HCl pH 7.5, 200 mM NaCl, 1 mM DTT for a total volume of 50 μL. Cytosine deamination to uracil percentage was measured with LC-MS/MS as described above.

### Synthesis of 5fC and 5caC modified DNA substrates

Oligonucleotides with 5fC and 5caC modified bases were synthesized in-house using standard phosphoramidite chemistry and supplier deprotection protocols (Glen Research, Sterling, VA). Control oligonucleotide (4CpG_contr) was purchased from Integrated DNA Technologies (IDT, Coralville, IA).

### Deaminase activity assay on 5fC and 5caC modified DNA

To test activity on 5fC and 5caC modified DNA, we designed a set of substrates (40 bp) containing 4 modified cytosines (Supplementary Table S4). The set of oligonucleotides included preferable deamination sites for 11 DNA deaminases from 5 representative clades. In each reaction, we mixed each modified oligonucleotide (dcaC or dfC) with the control oligonucleotide (C only) in a ratio of 1:1 (800 ng+800 ng) to monitor deamination of cytosine to uracil. After incubation for 5 h at 37°C in - reaction buffer containing 50 mM Bis-Tris pH 6.0, 0.1% Triton X-100 with different DNA deaminases, DNA was purified using Monarch PCR and DNA Cleanup kit, digested to nucleosides with the Nucleoside Digestion Mix (NEB, Ipswich MA) and the reaction products were quantified with LC-MS/MS.

### Deamination activity assay on RNA substrate

To test the activity of in-vitro expressed deaminases on RNA, 2 μL of freshly made PURExpress sample was mixed with 200 ng of heavy isotope labeled Fluc mRNA into buffer containing 50 mM Bis-Tris pH 6.0, 0.1% Triton X-100 and incubated for 1 h at 37°C for a total volume of 50 μL. Then, deaminase treated RNA was purified with NEBNext sample purification beads (E7104S, NEB, Ipswich, MA) using 1.2x beads to sample volume ratio, eluted in 40 μL nuclease free water. RNA sample was filtrated using a 0.22 µm centrifugal filter (UFC30GV00, MilliporeSigma, Darmstadt, Germany) to ensure all the beads were removed from the sample. Purified and filtrated RNA was then digested to single nucleoside by Nucleoside Digestion Mix (M0649S, NEB, Ipswich, MA). LC-MS/MS analysis was performed by injecting digested stable isotope labeled RNAs on an Agilent 1290 Infinity II UHPLC equipped with a G7117A diode array detector and a 6495C triple quadrupole mass detector operating in the positive electrospray ionization mode (+ESI). UHPLC was carried out on a Waters XSelect HSS T3 XP column (2.1 × 100 mm, 2.5 µm) with a gradient mobile phase consisting of methanol and 10 mM aqueous ammonium acetate (pH 4.5). MS data acquisition was performed in the dynamic multiple reaction monitoring (DMRM) mode.

Each nucleoside was identified in the extracted chromatogram associated with its specific MS/MS transition: rC* [M+H]+ at m/z 256.1→119.1; rU* [M+H]+ at m/z 256.1→119.1 (*: Stable isotope labeled nucleosides) External calibration curves with known amounts of the nucleosides were used to calculate their ratios within the samples analyzed. The conversion rate was calculated using the following formula:

labeled Uridine / (labeled Uridine + labeled C)

### DNA deamination NGS assay

50 ng of unmodified *E. coli* C2566 genomic DNA was combined with control DNAs with various DNA modification types (Supplementary Table S5) in 50 µL of 10 mM Tris pH 8.0.

Then the DNA was transferred to a Covaris microTUBE (Covaris, Woburn, MA) and sheared to 300 bp using the Covaris S2 instrument. 50 µL of sheared material was transferred to a PCR strip tube to begin library construction. NEBNext DNA Ultra II Reagents (NEB, Ipswich, MA) were used according to the manufacturer’s instructions for end repair, A-tailing, and adaptor ligation. The custom made Pyrollo-dC adaptor (NEB Organic Synthesis Division, Ipswich MA), where all dCs are replaced with Pyrollo-dC, was used: (ACACTCTTTCCCTACACGACGCTCTTCCGATC*T and [Phos]GATCGGAAGAGCACACGTCTGAACTCCAGTCA). The ligated samples were mixed with 110 µL of resuspended NEBNext Sample Purification Beads and cleaned up according to the manufacturer’s instructions. The library was eluted in 17 µL of water. For ssDNA libraries, prior to deamination the DNA was denatured by heating at 90°C for 10 minutes followed by cooling for 2 minutes on ice. The DNA was then deaminated in 50 mM Bis-Tris pH 6.0, 0.1% Triton X-100, using 1 µL of dsDNA deaminase synthesized as described above with an incubation time of 1 hour at 37°C. After deamination reaction, 1 µL of Thermolabile Proteinase K (NEB, Ipswich, MA) was added and incubated additional 30 min at 37°C followed by 10 min at 60°C. 5µM of NEBNext Unique Dual Index Primers and 25 µL NEBNext Q5U Master Mix (New England Biolabs, Ipswich, MA, USA) were added to the DNA and PCR amplified. The PCR reaction samples were mixed with 50 µL of resuspended NEBNext Sample Purification Beads and cleaned up according to the manufacturer’s instructions. The library was eluted in 15 µL of water. The libraries were analyzed and quantified by High sensitivity DNA analysis using a chip inserted into an Agilent Bioanalyzer 2100. The libraries were sequenced using the Illumina NextSeq and NovaSeq platforms. Paired-end sequencing of 150 cycles (2 x 75 bp) was performed for all the sequencing runs. Base calling and demultiplexing were carried out with the standard Illumina pipeline.

### Measure deaminase activity on N4mC

Deaminase activity on N4mC was measured from all the cacCgc sites in the genome of pRSSM1.PleII plasmid, of which the capitalized C is N4 methylated. To avoid the confounding effect of sequence selectivity, we only considered enzymes that have a minimum 10% activity on unmodified cacCgc sites in the *E. coli* DNA and calculated a relative deamination efficiency value, which is the percentage of one enzyme’s activity on N4mC compared to that on unmodified C in the same sequence context.

### SEM-seq library preparation of human genomic DNA

50 ng of human GM12878 genomic DNA spiked with 0.15 ng CpG-methylated pUC19, 1 ng unmethylated lambda DNA and 1 ng methylated XP12 DNA were sonicated with the Covaris S2 instrument, end repaired and ligated to the Pyrollo-dC adaptor as described above. The deamination of the resulting ligation product was performed with 1 µL of dsDNA deaminase (MsddA) in Low pH buffer (50 mM Bis-Tris pH 6.0, 0.1% Triton X-100) with the addition of 50mM NaCl and 1mM DTT in 20 µL total reaction volume for 3 h at 37 °C. The reaction was stopped by adding 1 µL of Thermolabile Proteinase K (NEB, Ipswich, MA) and incubating for 30 min at 37°C then an additional 10 min at 60°C. 15 µL of reaction without purification was directly used for PCR amplification (6 cycles) in 50 µL total volume and cleaned up as mentioned above. The whole genome libraries were sequenced on the Illumina NovaSeq 6000 platform (100 bp paired-end).

### Low input SEM-seq libraries

50 ng of human GM12878 genomic DNA spiked with 0.15 ng CpG-methylated pUC19, 1 ng unmethylated lambda DNA and 1 ng methylated XP12 DNA were sonicated with the Covaris S2 instrument. Sheared DNA was diluted to a concentration of 0.2 pg/µL, from which 50 µL (10 pg) of were used for end repair and adaptor ligation as described above. For adaptor ligation Pyrollo-dC adaptor was diluted 25 times.

The deamination of the resulting ligation product was performed with 1 µL of dsDNA deaminase (MsddA) in Low pH buffer (50 mM Bis-Tris pH 6.0, 0.1% Triton X-100) with the addition 15 ng of Lambda DNA (Promega, Fitchburg, WI) in 20 µL total reaction volume for 3 h at 37 °C. The reaction was stopped by adding 1 µL of Thermolabile Proteinase K (NEB, Ipswich, MA) and incubating for 30 min at 37°C then an additional 10 min at 60°C. 15 µL of reaction without purification was directly used for PCR amplification (12 and 14 cycles) in 50 µL total volume and cleaned up as mentioned above two times. The whole genome libraries were sequenced on the Illumina NovaSeq 6000 platform (100 bp paired-end).

### Data analysis

#### Read processing and mapping

Raw Illumina reads were first trimmed by the Trim Galore software (https://github.com/FelixKrueger/TrimGalore) to remove adapter sequences and low-quality bases from the 3’ end. Unpaired reads due to adapter/quality trimming were also removed during this process. The trimmed read sequences were C to T converted and were then mapped to the corresponding reference using the Bismark program^44^ with the default Bowtie2 settings^45^. For the NGS deamination assay, a composite reference was created by combining the complete genome sequences of *E.* coli C2566, phage lambda, phage XP12, phage T4, plasmid pRRSlac, and Adenovirus. For human libraries made by SEM-seq, EM-seq and WGBS, a composite reference included the human genome (GRCh38) and the complete sequences of unmethylated lambda and CpG-methylated pUC19 controls.

The first 5 bp at the 5’ end of R2 reads were removed to reduce end-repair errors and aligned read pairs that shared the same alignment ends were regarded as PCR duplicates and were discarded. For human libraries, aligned reads that contained excessive cytosines in non-CpG context (more than 4 in 100bp) were removed before calculating methylation level for individual sites because these reads likely contain high conversion errors.

The numbers of Ts (converted not methylated) and Cs (unconverted modified) of each covered cytosine position were then calculated from the remaining good quality alignments using Bismark methylation extractor.

Analysis of deamination efficiency and sequence preference from the NGS data Deamination events were inferred from C to T changes at each covered cytosine position in the genomes based on Bismark methylation extractor output. The 20 bp flanking sequences (10 bp upstream and 10 bp downstream) were extracted from all the covered cytosines that had at least 10X read coverage from the individual genomes and divided the cytosines sites into different groups based on their deamination rates (e.g., >=90%, >=50%). Flanking sequences of each cytosine group were used to make sequence logo using WebLogo 3^22^ to infer deamination sequence preference.

#### Determination of methylated CpG sites in human genome by Binomial correction

For all samples except the 10 pg SEM-seq libraries, to avoid false identification of methylated positions due to the incomplete conversion of unmodified cytosines, we used Binomial statistics with Benjamini-Hochberg correction to determine the methylated CpG sites that have significantly higher methylation levels compared with the background. The P-value of the one-sided binomial test (𝐻_0_: 𝑃 ≤ 𝑃_0_, 𝐻_𝑎_: 𝑃 > 𝑃_0_) can be calculated as 𝑃 = 1 − 𝐹(𝑘; 𝑛, 𝑃_0_), where 𝐹 is the cumulative distribution function of the binomial distribution, 𝑘 is the number of unmethylated cytosine, 𝑛 is the coverage (number of methylated and unmethylated cytosine), 𝑃 is the sample methylation level for each region, and 𝑃_0_is the background methylation level which is the non-conversion rate estimated from unmethylated lambda genome. The P-value was further adjusted using Benjamini-Hochberg method. Only the CpG positions with coverage above a certain cutoff and FDR (false discovery rate) < 0.05 were defined as methylated CpG sites. For a single replicate and replicates combined data, we require a minimum of 3X and 5X coverage, respectively.

#### Genome-wide coverage analysis

We combined the filtered bam files from the two technical replicates to investigate the sequencing coverage across the genome. The GC content bias statistics were calculated using Picard (version 2.26.11) CollectGcBiasMetrics (http://broadinstitute.github.io/picard) and was plotted using a custom python script.

#### Determination of coverage around CpG islands and transcription start sites (TSS)

We combined the filtered bam files from the two technical replicates to examine the sequencing coverage of different methods around the CpG islands and transcription start sites. The reference bed files of human CpG islands and TSS were downloaded from UCSC Table Browser^46^. and The coverage was computed using bamCoverage from deepTools (version 3.3.1)^47^ with the following parameters: -- binSize 50 --normalizeUsing RPGC --effectiveGenomeSize 2913022398 –ignoreDuplicates. Then the coverage was calculated for per genome regions using computeMatrix from deepTools with -a 1000 -b 1000; and scale-regions and reference-point option were used for CpG islands and TSS, respectively.

#### Determination of methylation status of CpG sites at epigenomic regions

We studied the methylation status of defined CpG sites at key epigenetic regions. We used GM12878 ChIP-seq data sets (processed hg38 bed narrowPeak files) from ENCODE portal^23^: ENCFF023LTU (H3K27ac), ENCFF188SZS (H3K4me3), and ENCFF796WRU (CTCF). The methylation of CpG sites around these epigenetic regions was calculated using computeMatrix reference-point from deepTools with the following parameters: -a 1000 -b 1000 --skipZeros --referencePoint center.

## Supplementary information

Supplementary Table 1: Supplementary_Table_1.xlsx

Supplementary Figures 1-16, and Supplementary tables S2-S6: Supplementary_Figures_and_tables.pdf

## Data Availability

NGS substrate specificity libraries and SEM-seq libraries are available at NCBI GEO accession: GSE233932

## Acknowledgements

We thank Yan-Jiun Lee and Peter Weigele for providing XP12 phage DNA, Dr. Lidija Truncaite, Vilnius University Life Sciences Center, Vilnius Lithuania for fully C-hydroxymethylated T4147 and fully C-hydroxy-methyl-beta-glucosylated T4 alpha-glucosyl-transferase knockout (AGT-) genomic DNAs, Dr. Chen Song for cfDNA. pRSSM1.PleII plasmid (containing N4mC at cacCgc sites was obtained from Dr. Iain Murray and Richard J. Roberts). We thank Dora Posfai, Laurie Mazzola, Danielle Fuchs, Bell Harold, and Kristen Augulewicz from the Sequencing Core at New England for performing Illumina sequencing. We are also grateful for the critical readings of our manuscript by Richard Roberts, Ira Schildkraut, Sriharsa Pradhan, and colleagues at NEB. This work was supported by New England Biolabs Inc.

