## supplemental material for "Discovery of novel DNA cytosine deaminase activities enables a nondestructive single-enzyme methylation sequencing method for base resolution high-coverage methylome mapping of cell-free and ultra-low input DNA"

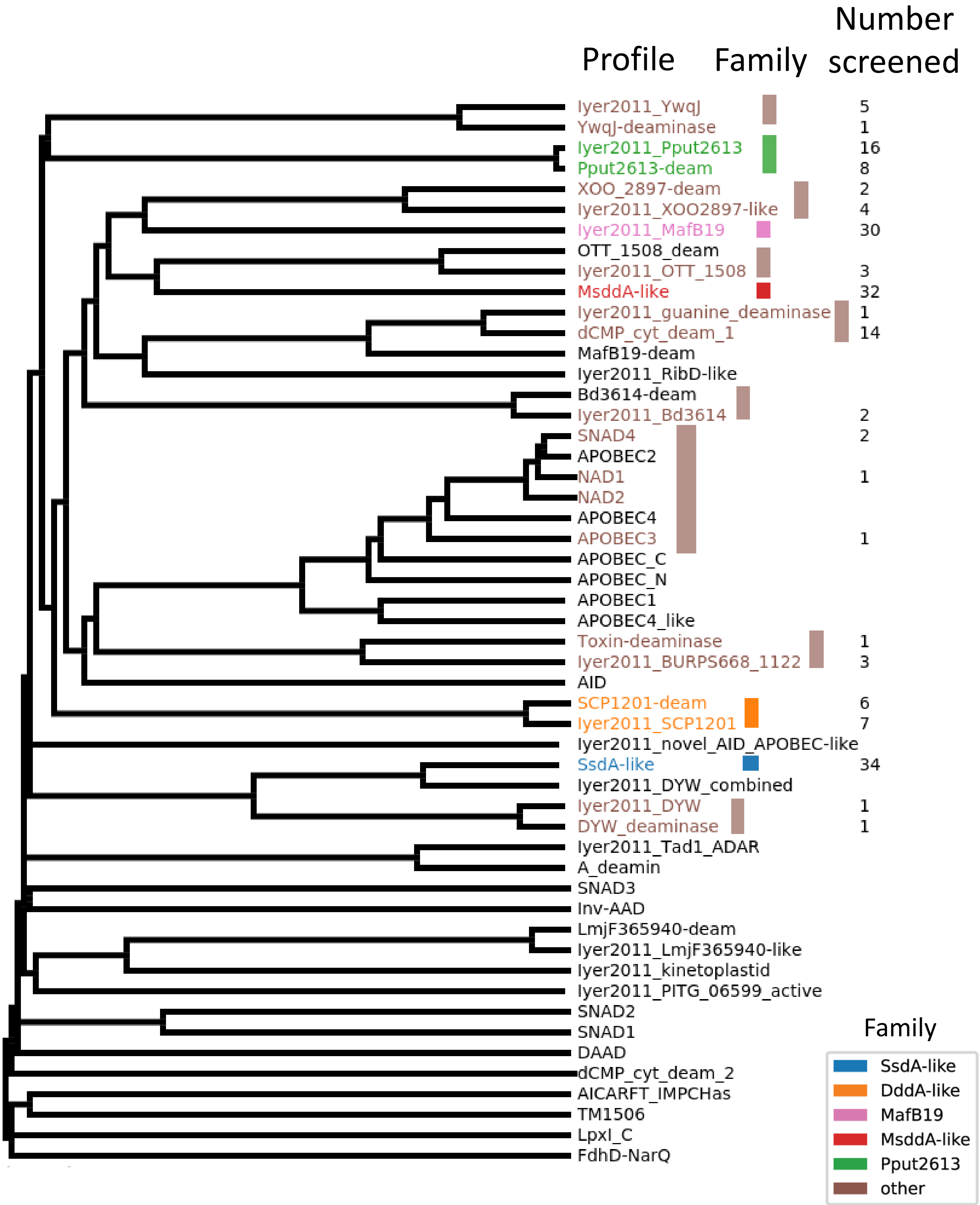


**Supplementary Figure 1. UPGMA tree of hmm profiles from the Pfam CDA clan (CL0109), Iyer et al. (2011), and Zhang et al. (2012).** Colored bars show groupings of the profiles into families. Numbers of screened enzymes with the highest score to each profile is indicated. Almost all active enzymes in our screen came from the five families highlighted with their own color.

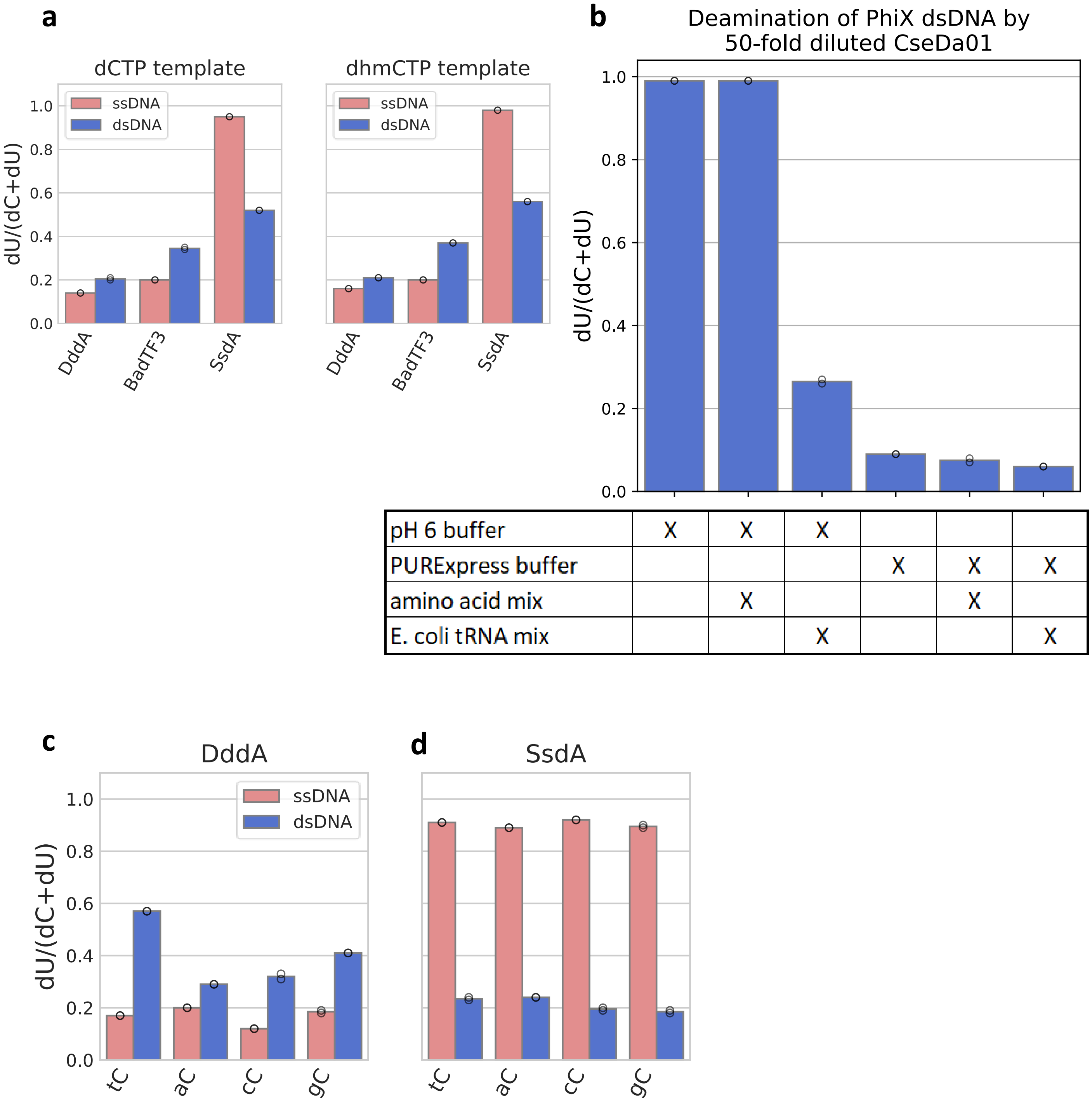


| | | | | | | |
|---|---|---|---|---|---|---|
| pH 6 buffer | X | X | X | | | |
| PURExpress buffer | | | | X | X | X |
| amino acid mix | | X | | | X | |
| E. coli tRNA mix | | | X | | | X |

**Supplementary Figure 2. Validation of NEB PURExpress system for expression of active bacterial DNA deaminases.** a) Comparison of deamination efficiency in pH 6 buffer, measured by the LC-MS assay, from enzyme produced in PURExpress reactions using transcription template containing unmodified dCTP, or dhmCTP. b) Deamination of PhiX dsDNA by 50-fold diluted CseDa01 (2 µL taken from a dilution of 2 µL in-vitro translation product into 98 µL water) in various buffers, measured by LC-MS assay, two technical replicates of each condition. PURExpress buffer is composed of PURExpress Solution A (-aa, tRNA) and Solution B from the PURExpress® Δ (aa, tRNA) Kit (New England Biolabs catalog #E6840S). Amino acid mix and E. coli tRNA mix are from the same kit (#E6840S). c) DddA and d) SsdA deamination efficiency on single-stranded or double-stranded 44 bp oligo substrates with cytidines in the contexts TC, AC, CC, and GC in pH 7.5 buffer. Two technical replicates were run of each condition in this figure.

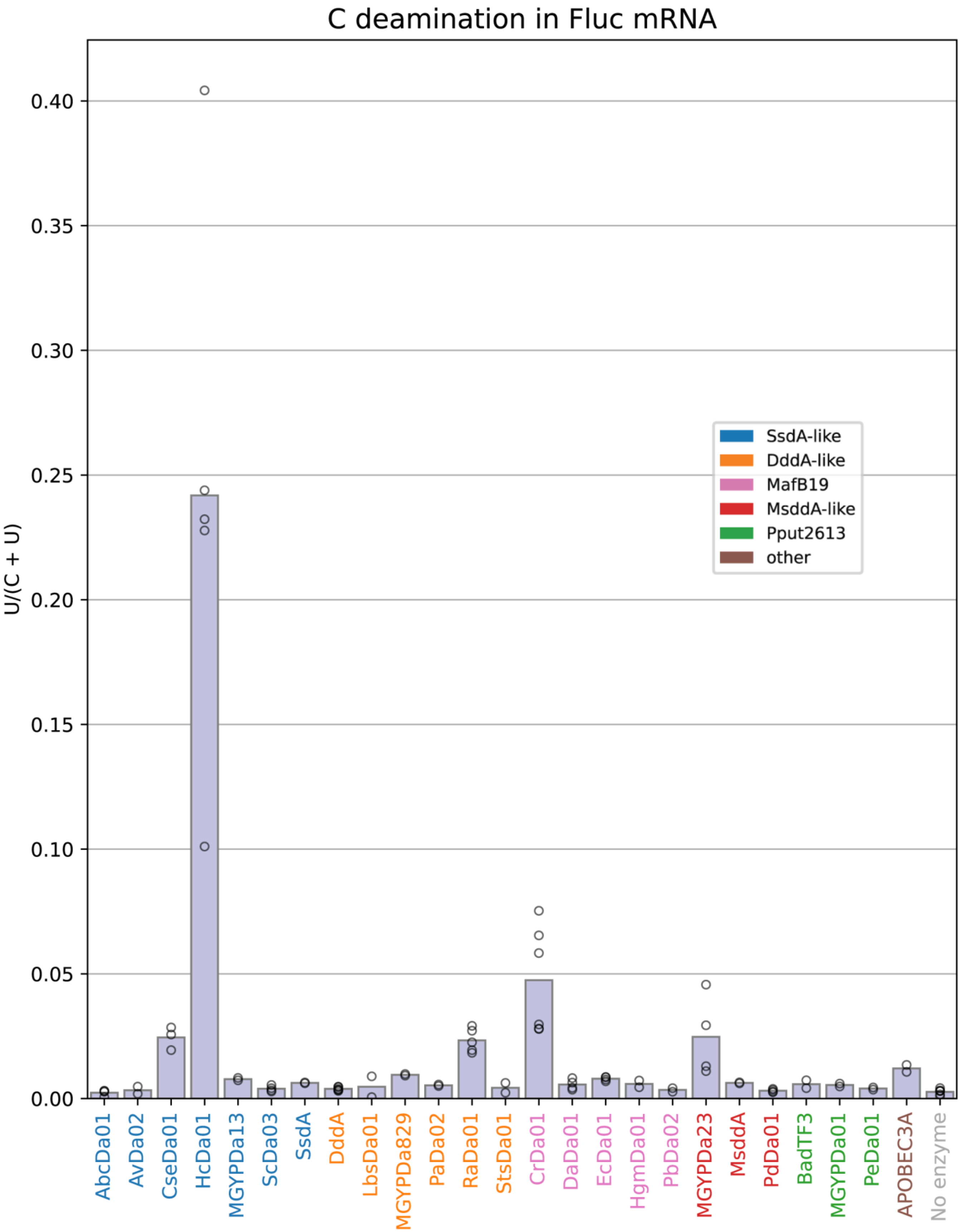


**Supplementary Figure 3. C deamination activity on Fluc mRNA substrate.**

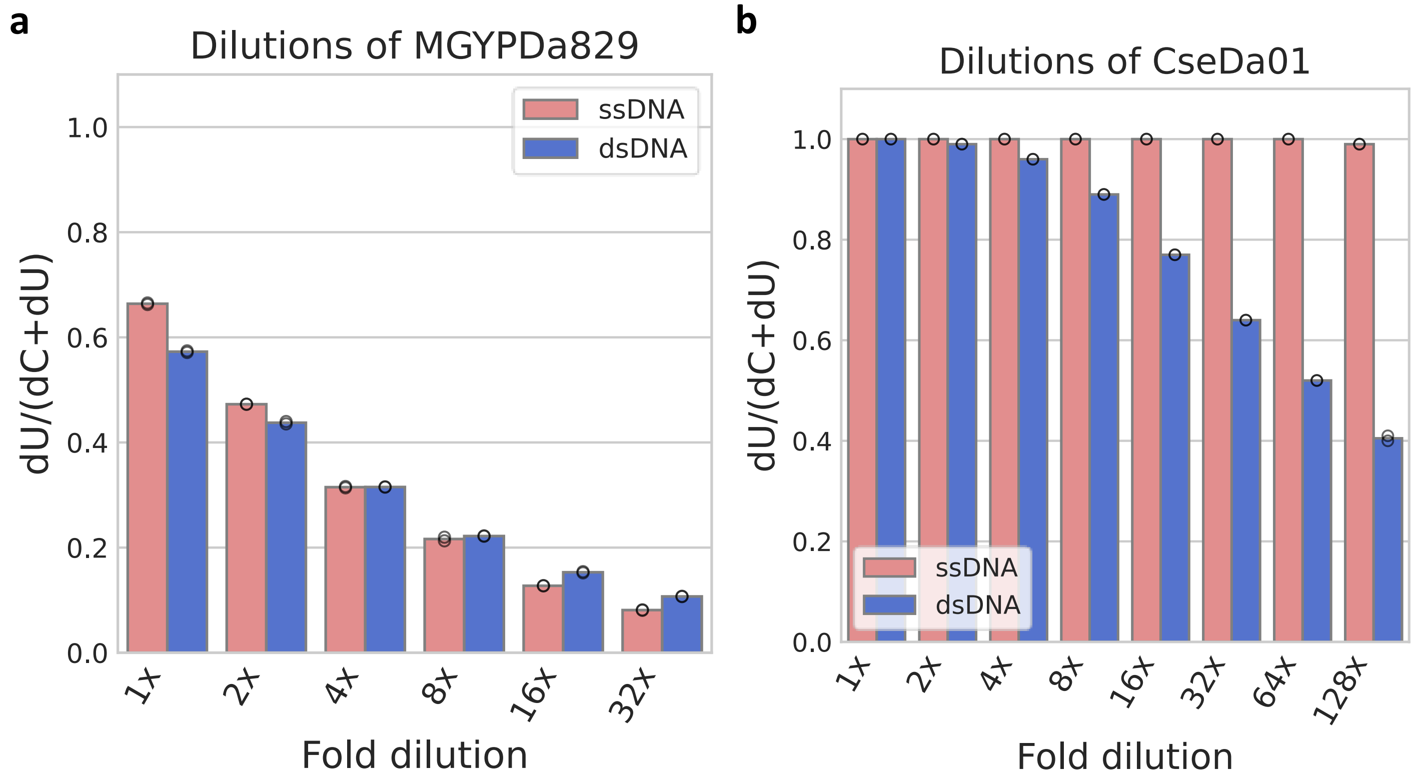


**Supplementary Figure 4. Effects of dilution on preference for ssDNA or dsDNA** of a) MGYPDa829 and b) CseDa01 as measured by LC-MS assay.

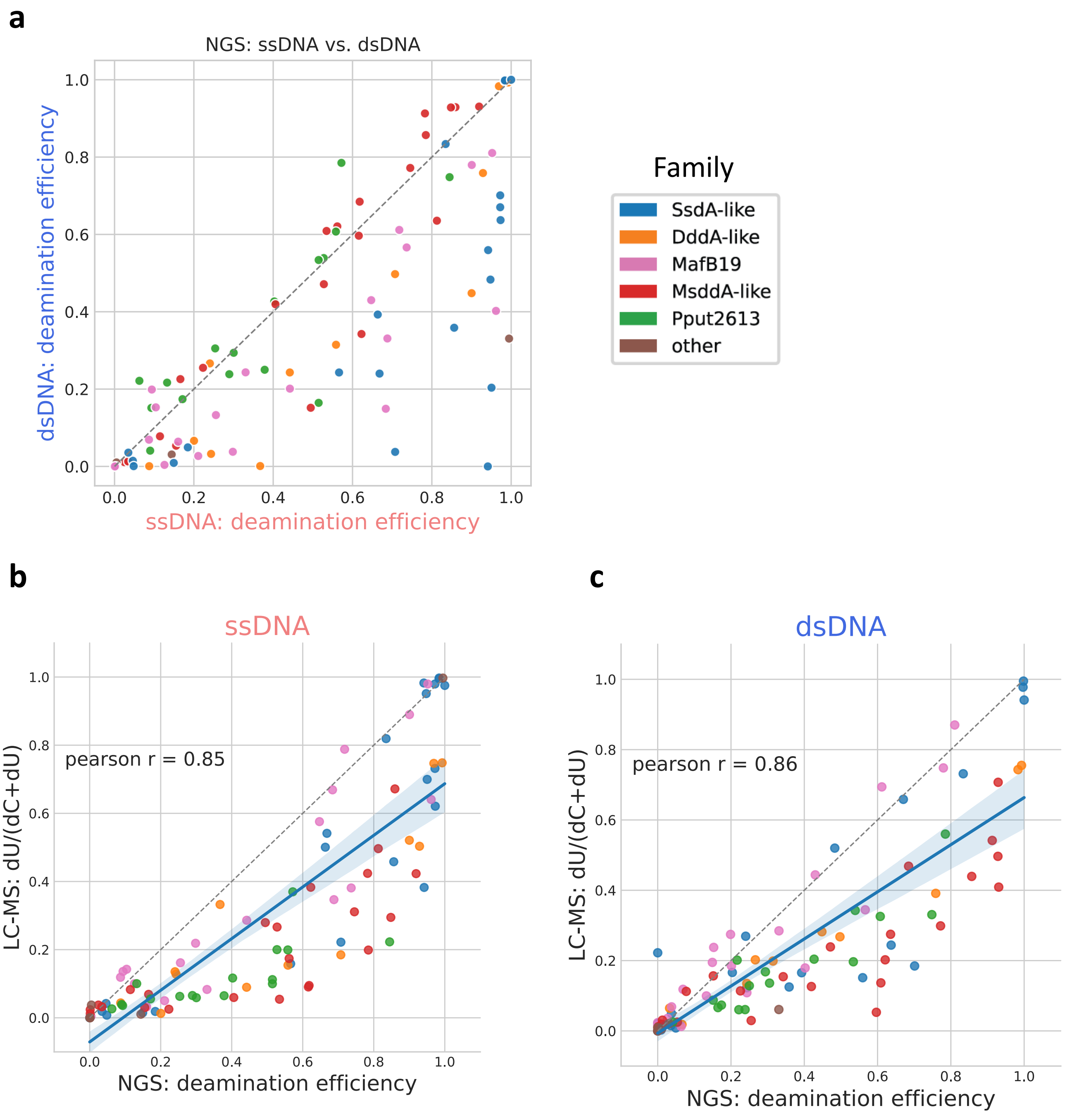


**Supplementary Figure 5. Comparison of ssDNA and dsDNA deamination efficiency** as measured by a) the NGS assay, and as compared between the NGS assay and LC-MS assay in b) ssDNA and c) dsDNA. Dashed grey line marks the diagonal (x=y). Blue lines are linear regressions with a 95% confidence interval.

a.

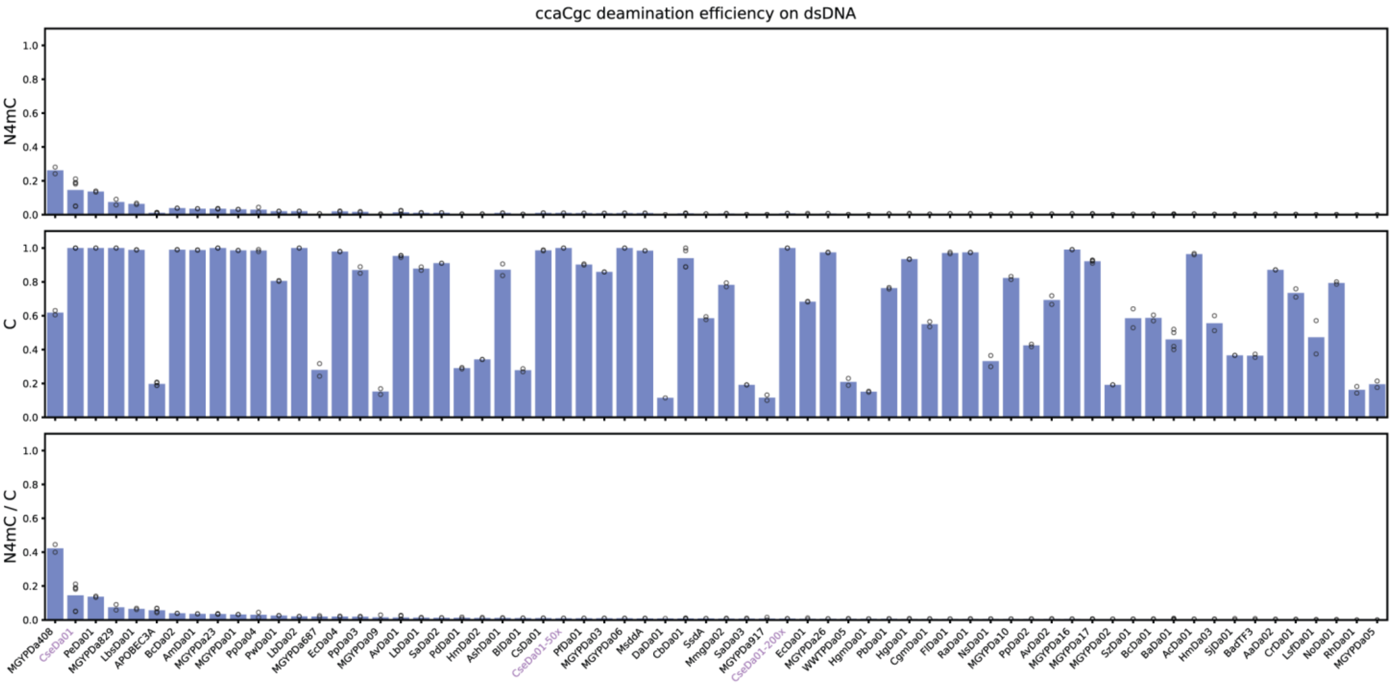


b.

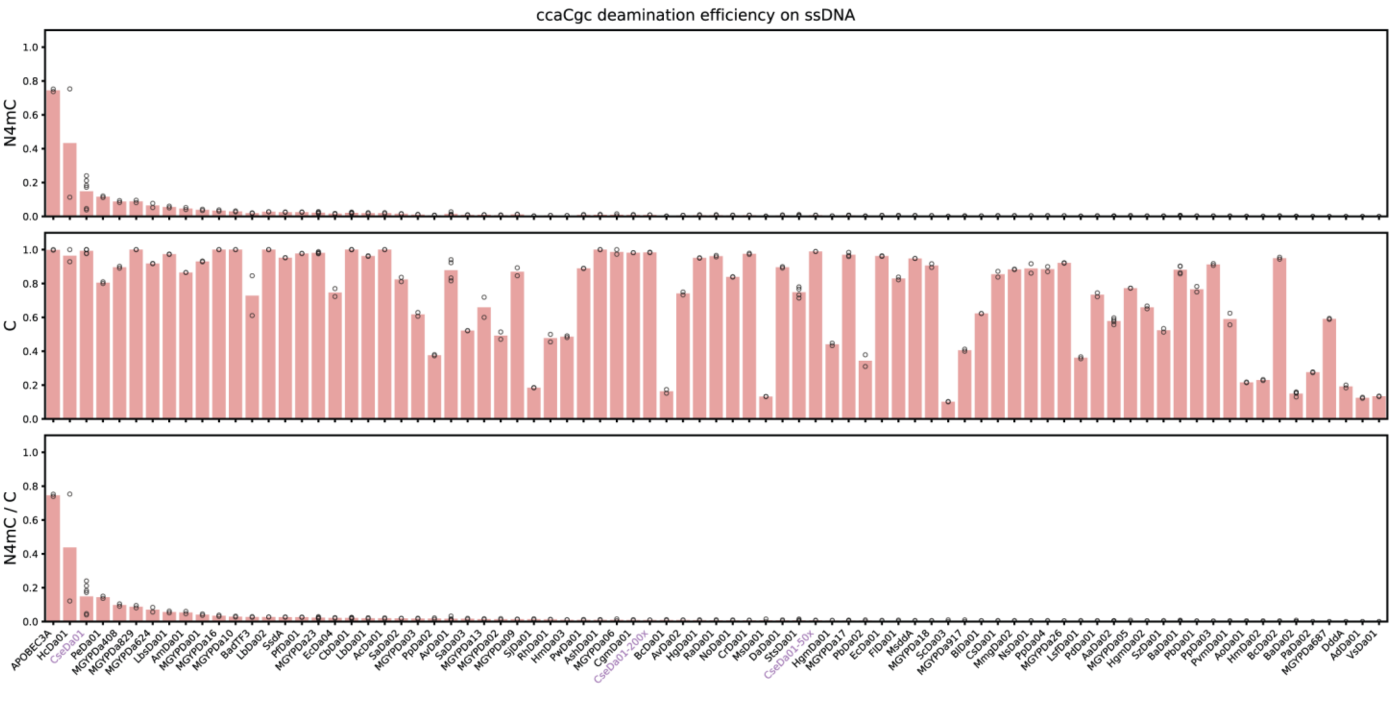


**Supplementary Figure 6. Deamination efficiency on ccaCgc measured by NGS assay.** a) dsDNA, b) ssDNA. Three dilutions of CseDa01 were assayed (violet labels), full strength, 50-fold dilution, and 200-fold dilution.

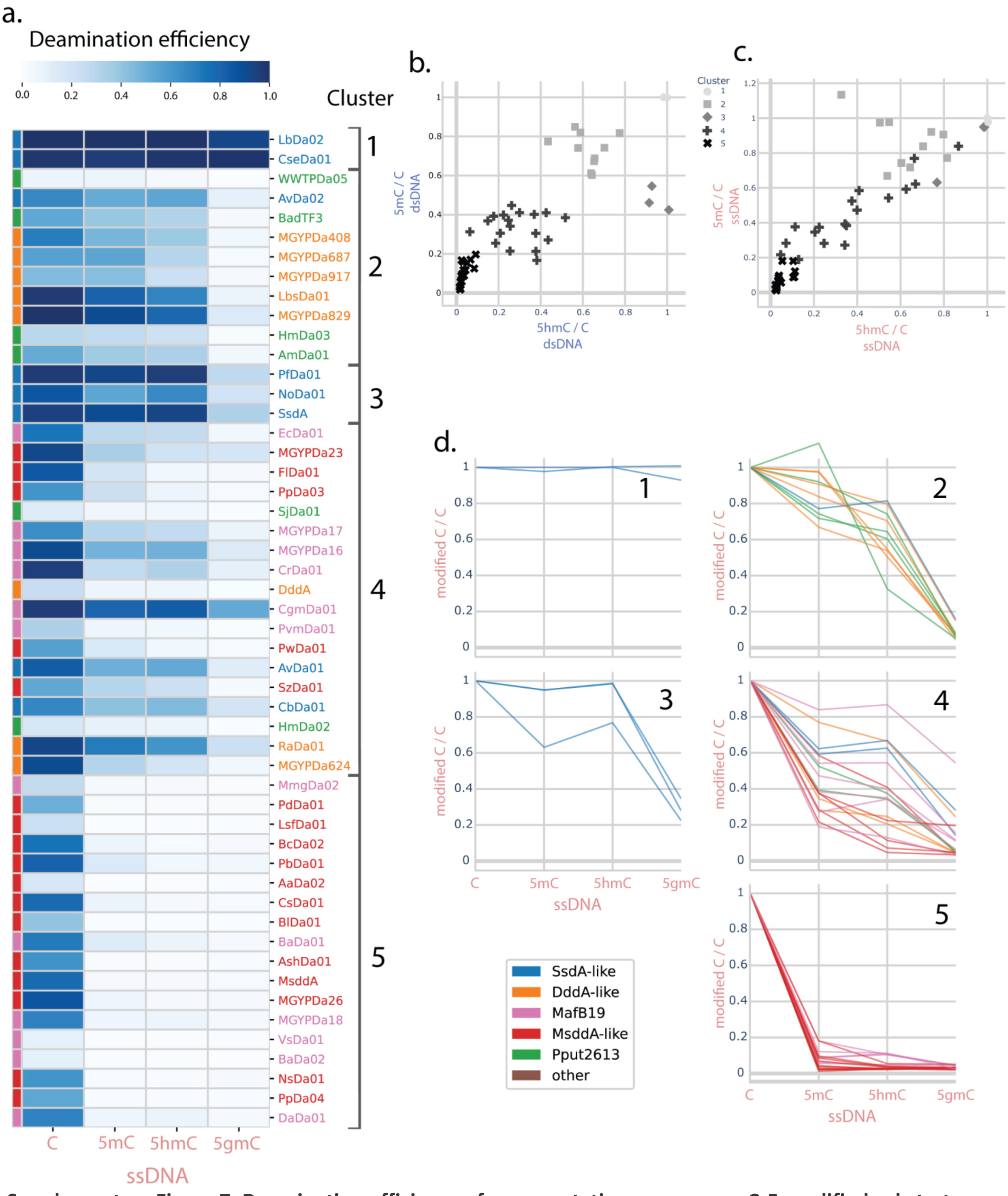


**Supplementary Figure 7. Deamination efficiency of representative enzymes on C-5 modified substrates in ssDNA measured by NGS assay.** a) Deamination efficiency on C, 5mC, 5hmC, and 5ghmC, clustering is based on dsDNA and is the same as in main text Fig. 2. b) Deamination efficiency on dsDNA 5mC and 5hmC divided by deamination efficiency on C. c) Deamination efficiency on ssDNA 5mC and 5hmC divided by deamination efficiency on C. d) Deamination efficiency on ssDNA C, 5mC, 5hmC, and 5ghmC divided by deamination efficiency on C.

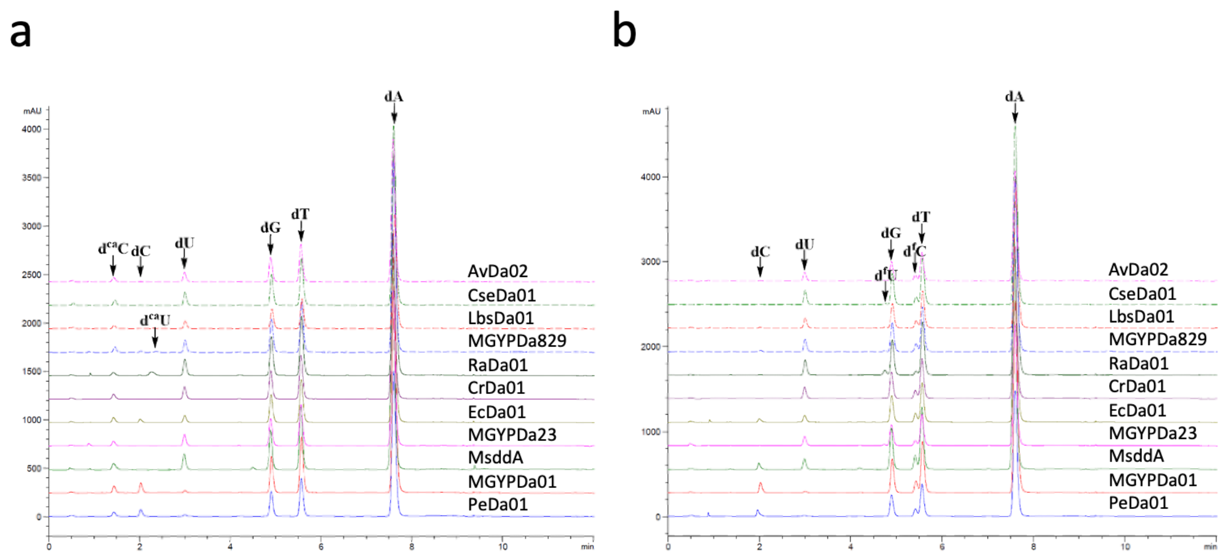


**Supplementary Figure 8. LC-MS total ion chromatograms of deamination assays on d5caC (a) and d5fC (b) ssDNA oligonucleotides.**

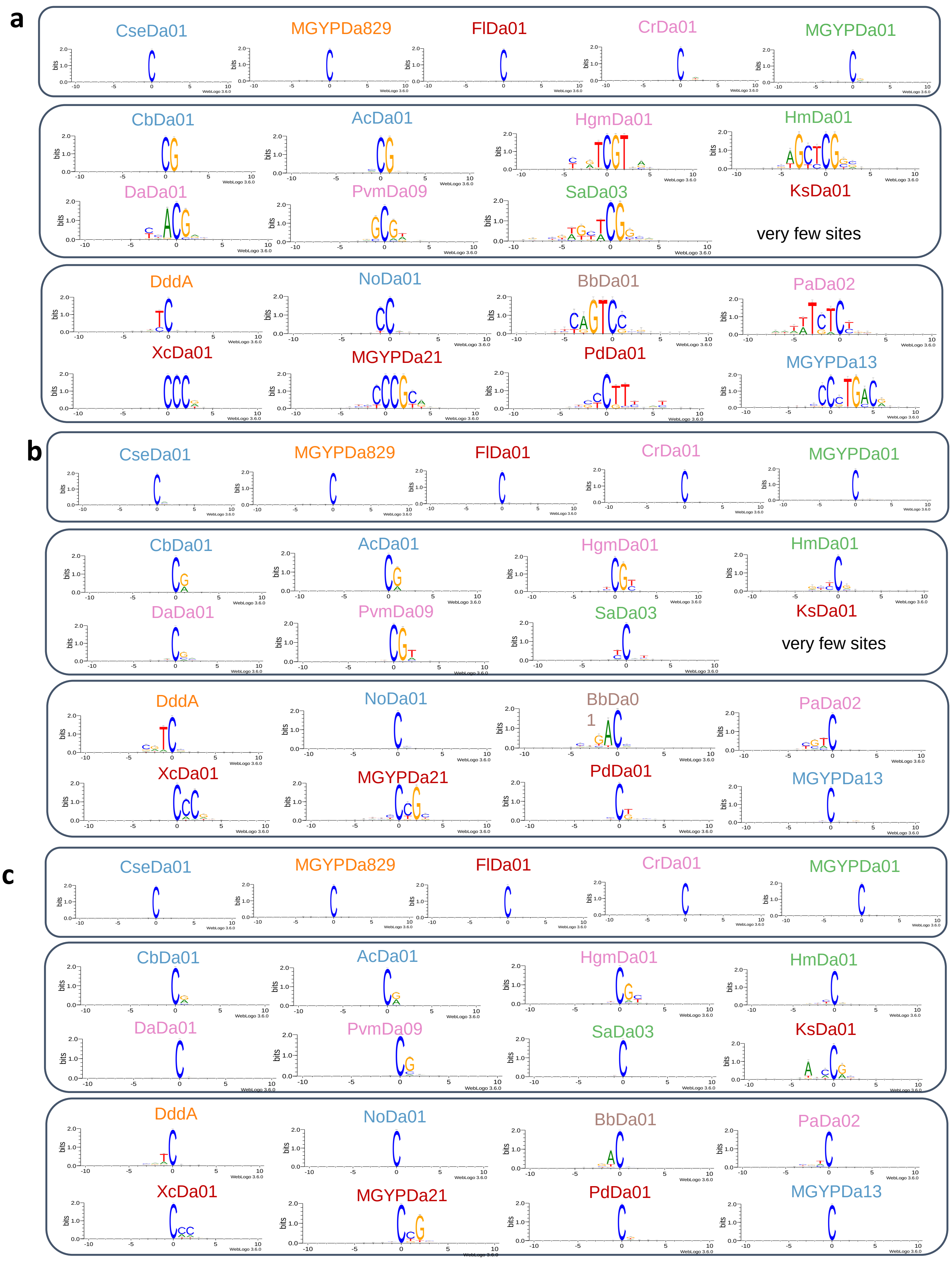


**Supplementary Figure 9. Diverse sequence preferences of deaminases.** a) Logos of sites with >=90% deamination efficiency in unmodified dsDNA. b) Logos of sites with >=90% deamination efficiency in unmodified ssDNA. c) Logos of sites with >=50% deamination efficiency in unmodified ssDNA substrates.

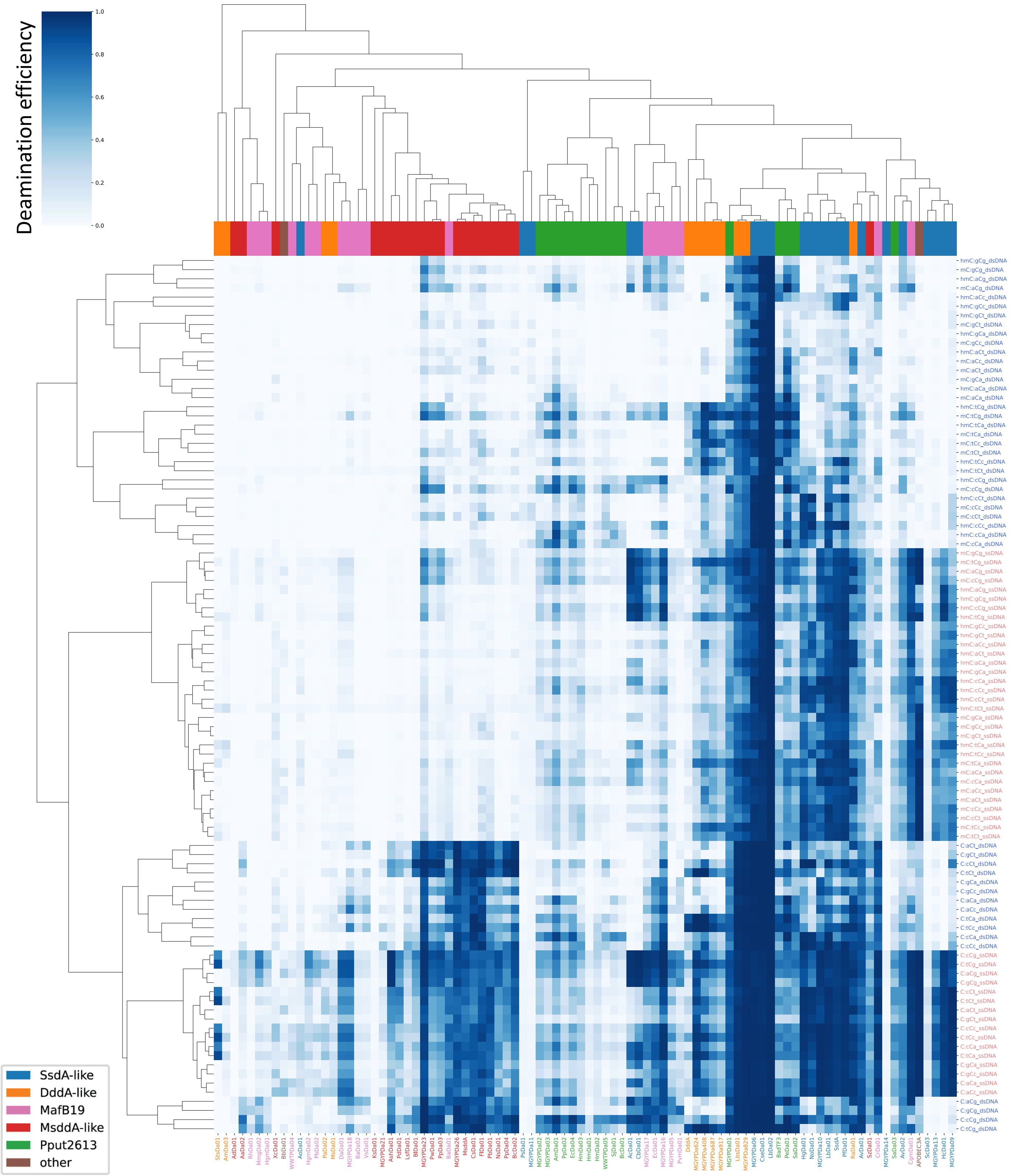


**Supplementary Figure 10. Deamination efficiency of representative deaminases on nCn contexts of modified and unmodified dsDNA and ssDNA.** Rows and columns are sorted based on average linkage clustering of cosine distances.

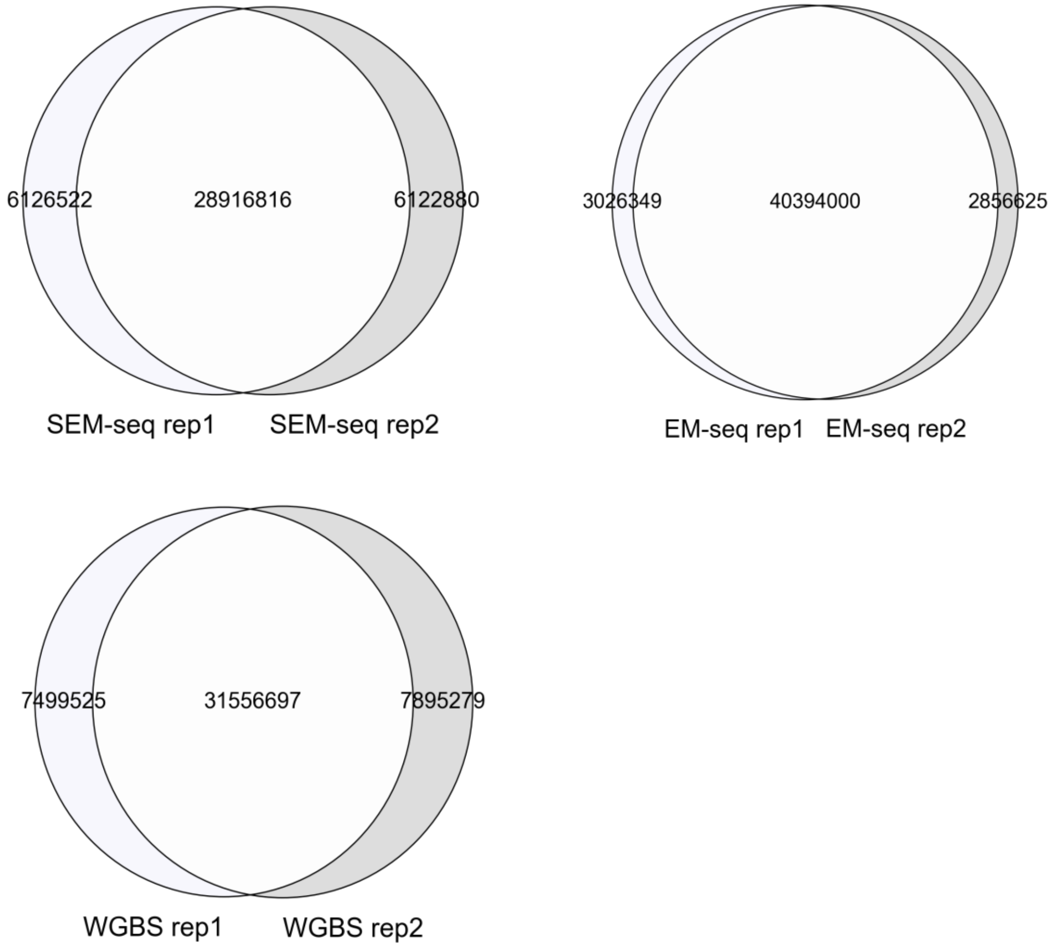


**Supplementary Figure 11. Comparison of methylated CpG sites between the 2 technical replicates of SEM-seq (50 ng), EM-seq (50 ng), and WGBS.**

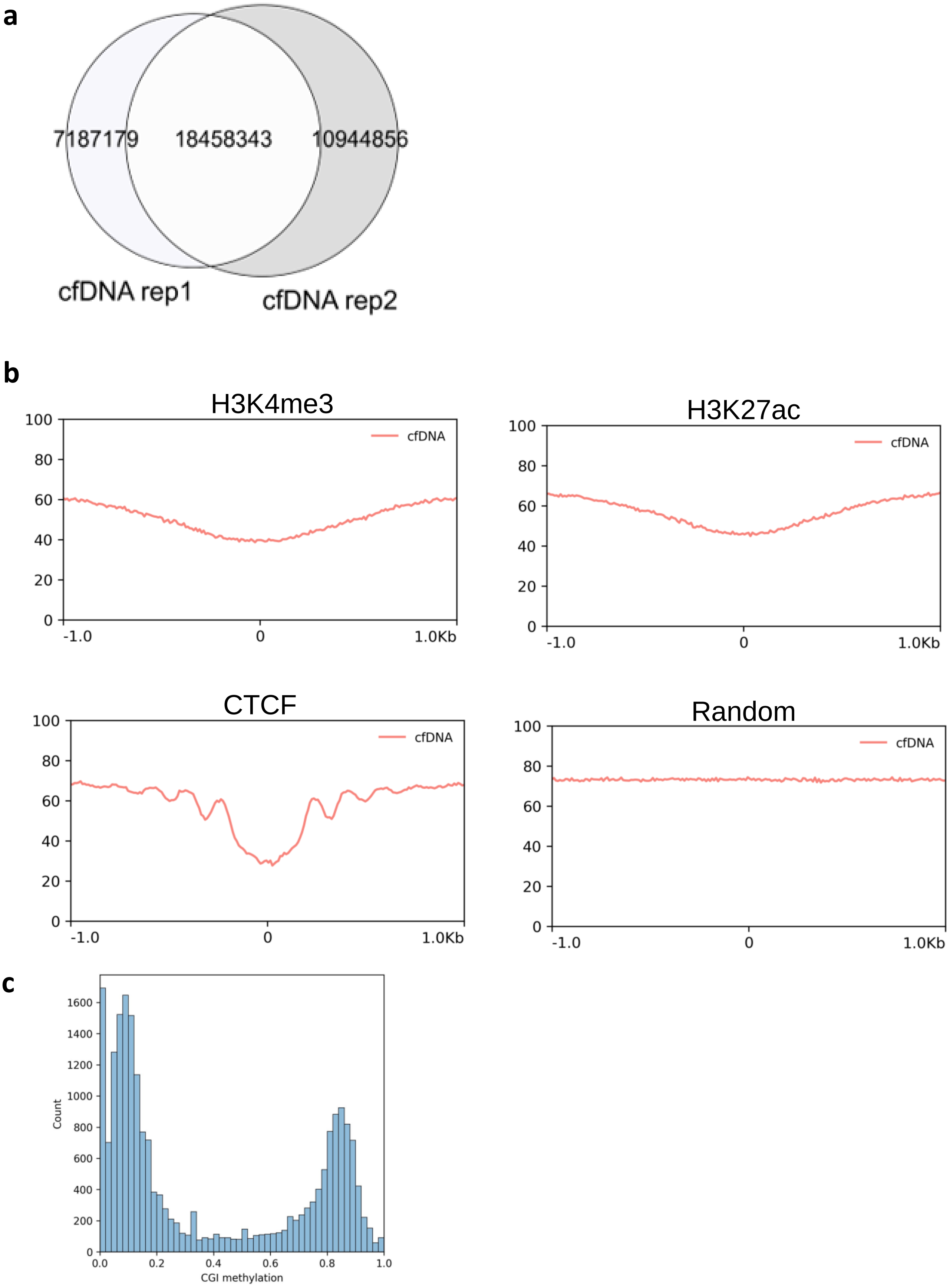


**Supplementary Figure 12. CfDNA SEM-seq library.** a) Comparison of methylated CpG sites between the 2 technical replicates. b) CpG methylation profiles in neighborhoods around specific genomic features. c) Distribution of cfDNA_02 (replicate 2) CpG island methylation quantified by SEM-seq.

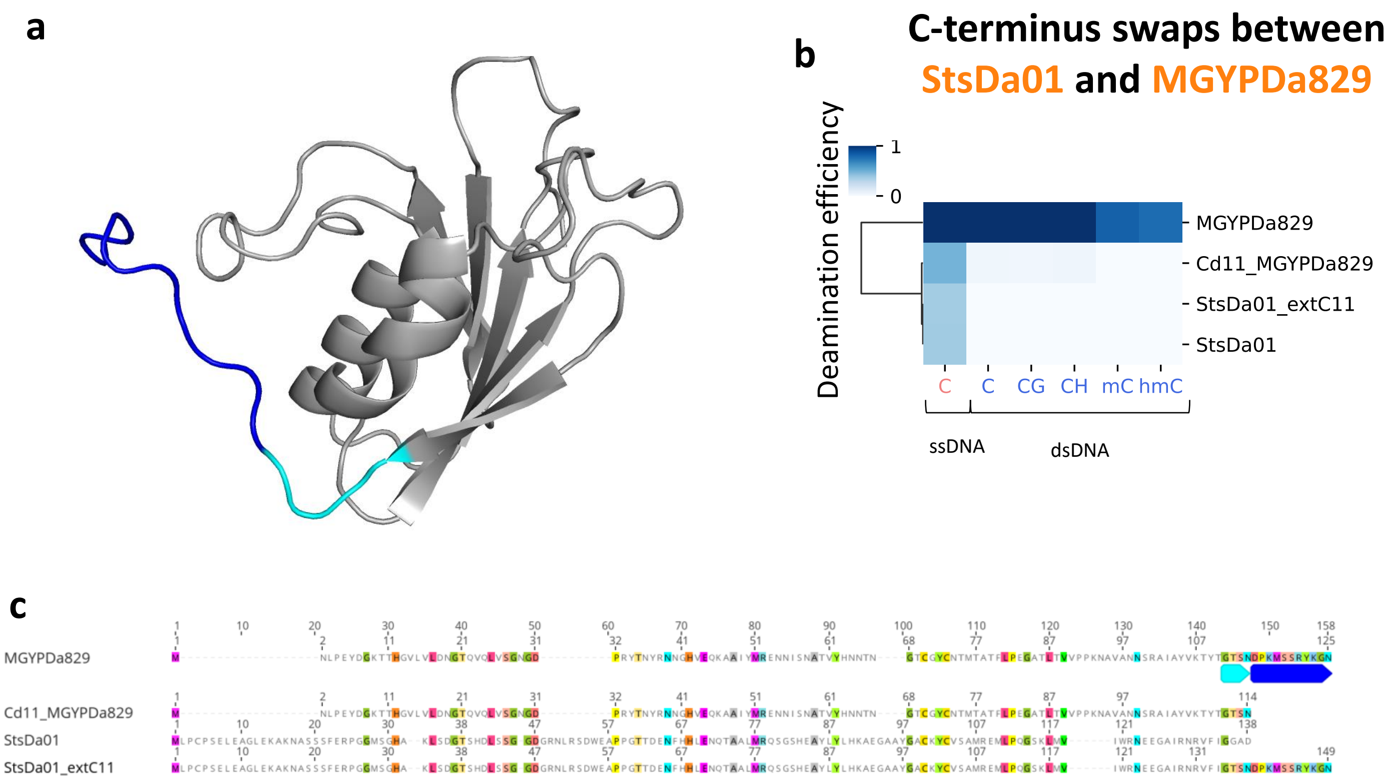


**Supplementary Figure 13. Effects of C-terminal swaps between MGYPDa829 and StsDa01.** a) MGYPDa829 structure predicted by AlphaFold2, shades of blue mark deleted or swapped regions. b) Heatmap of deamination efficiency on various substrates, measured by NGS assay. Clustering is cosine distance, average linkage. Names of the enzymes are colored orange to indicate that they are from the DddA-like family. c) Multiple sequence alignment of MGYPDa829, StsDa01, and their chimeras. Blue boxes correspond to the regions of the structure with the same colors.

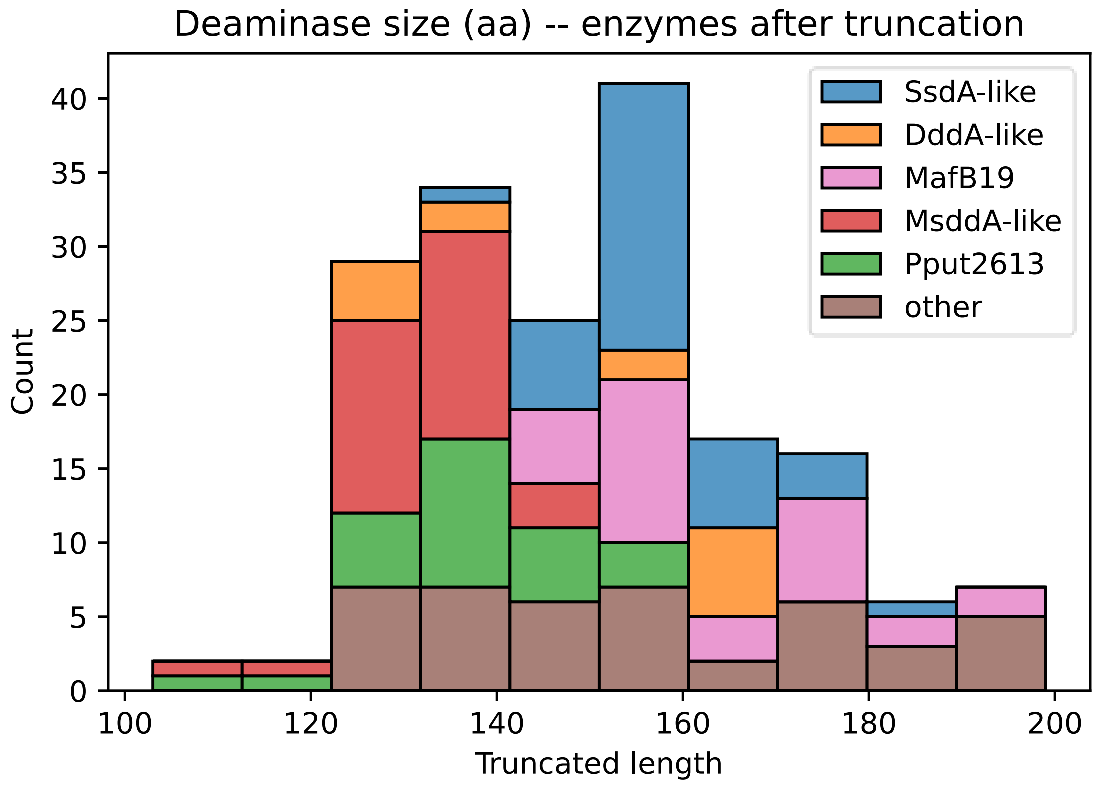


**Supplementary Figure 14. Length distribution of screened deaminases.**

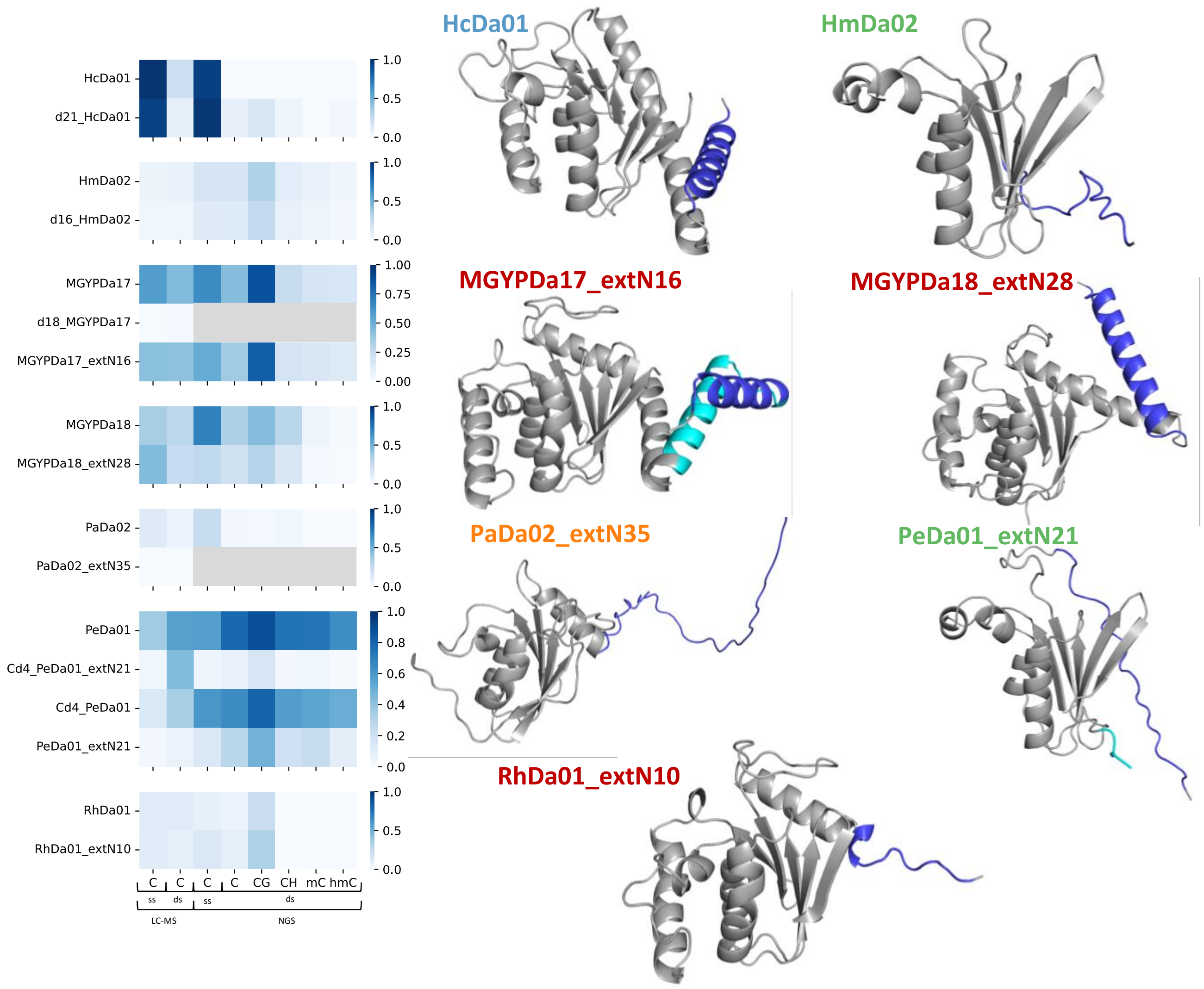


**Supplementary Figure 15. Activities of alternative truncations of various deaminases.** Structures are AlphaFold2 predictions. Colored sections of predicted structures indicate sections added or removed from the core (named with no prefix or suffix) sequence. In some cases where low or no activity was seen in the LC-MS assay, NGS assays were not run, and are marked-out in grey.

### Supplementary Table S2. Gblock amplification Primers

| NAME | SEQUENCE |
|---|---|
| T7_FW | GCGAATTAATACGACTCACTATAGGG |
| T7_R | CCAAATAAACCCCTCCGTTTAGA |

### Supplementary Table S3. Oligonucleotide sequences for DddA and SsdA

| NAME | SEQUENCE |
|---|---|
| DddA_TC_Top | AAAAAAAAAAAAAAAATCAAAAAAAAAAAAAAAA |
| DddA_AC_Top | AAAAAAAAAAAAAAAAACAAAAAAAAAAAAAAAA |
| DddA_CC_Top | AAAAAAAAAAAAAAAACCAAAAAAAAAAAAAAAA |
| DddA_GC_Top | AAAAAAAAAAAAAAAAGCAAAAAAAAAAAAAAAA |
| DddA_TC_Bottom | TTTTTTTTTTTTTTTTGATTTTTTTTTTTTTTTT |
| DddA_AC_Bottom | TTTTTTTTTTTTTTTTGTTTTTTTTTTTTTTTTT |
| DddA_CC_Bottom | TTTTTTTTTTTTTTTTGGTTTTTTTTTTTTTTTT |
| DddA_GC_Bottom | TTTTTTTTTTTTTTTTGCTTTTTTTTTTTTTTTT |

### Supplementary Table S4. Oligonucleotide sequences for 5fC and 5caC

| NAME | SEQUENCE |
|---|---|
| 4CpG_5fC | AAATTTAATTATAAAA(5fC)GAT(5fC)GA(5fC)GA(5fC)GAAATAATAAAAA |
| 4CpG_5caC | AAATTTAATTATAAAA(5caC)GAT(5caC)GA(5caC)GA(5caC)GAAATAATAAAA |
| 4CpG_contr | AAATTTAATTATAAAACGATCGACGACGAAATAATAAAAA |

**Supplementary Table S5. Substrates in NGS activity assay**

| DNA | MODIFICATION | DNA amount (ng) |
|---|---|---|
| E. coli C2566 | C | 46.8 |
| Lambda phage, dcm- | C | 1 |
| XP12 phage | 5mC | 1 |
| 1783 bp PCR fragment of Adenovirus genome (below) amplified with d5hmCTP using primers in Table S5 | 5hmC | 0.1* |
| Fully C-hydroxymethylated T4147 | 5hmC | 1* |
| T4 phage, AGT- | 5gmC | 1* |
| pRSSM1.PleII | N4mC | 0.1 |

*Only one of the 5hmC substrates was used in each assay, with some using T4147 and others using the amplified adenovirus fragment. T4147 and T4 phage, AGT- were never used in the same libraries.

**Supplementary Table S6. Adenovirus 5hmC substrate primers**

| NAME | SEQUENCE |
|---|---|
| Adeno_5hmC_F | CGAAGCTATGTCCAAGCGC |
| Adeno_5hmC_R | CTGTTAATCTTATTTTGCACTGCCTG |

>Adenovirus 5hmC 1783 bp PCR fragment

```
CGAAGCTATGTCCAAGCGCAAAATCAAAGAAGAGATGCTCCAGGTCATCGCGCCGGAGATCTATGGCCCCCCGAAGA
AGGAAGAGCAGGATTACAAGCCCCGAAAGCTAAAGCGGGTCAAAAAGAAAAAGAAAGATGATGATGATGATGAACTT
GACGACGAGGTGGAACTGCTGCACGCAACCGCGCCCAGGCGGCGGGTACAGTGGAAAGGTCGACGCGTAAGACGTGT
TTTGCGACCCGGCACCACCGTAGTTTTTACGCCCGGTGAGCGCTCCACCCGCACCTACAAGCGCGTGTATGATGAGG
TGTACGGCGACGAGGACCTGCTTGAGCAGGCCAACGAGCGCCTCGGGGAGTTTGCCTACGGAAAGCGGCATAAGGAC
ATGTTGGCGTTGCCGCTGGACGAGGGCAACCCAACACCTAGCCTAAAGCCCGTGACACTGCAGCAGGTGCTGCCCAC
GCTTGCACCGTCCGAAGAAAAGCGCGGCCTAAAGCGCGAGTCTGGTGACTTGGCACCCACCGTGCAGCTGATGGTAC
CCAAGCGCCAGCGACTGGAAGATGTCTTGGAAAAAATGACCGTGGAGCCTGGGCTGGAGCCCGAGGTCCGCGTGCGG
CCAATCAAGCAGGTGGCACCGGGACTGGGCGTGCAGACCGTGGACGTTCAGATACCCACCACCAGTAGCACTAGTAT
TGCCACTGCCACAGAGGGCATGGAGACACAAACGTCCCCGGTTGCCTCGGCGGTGGCAGATGCCGCGGTGCAGGCGG
CCGCTGCGGCCGCGTCCAAAACCTCTACGGAGGTGCAAACGGACCCGTGGATGTTTCGCGTTTCAGCCCCCCGGCGC
CCGCGCCGTTCCAGGAAGTACGGCACCGCCAGCGCACTACTGCCCGAATATGCCCTACATCCTTCCATCGCGCCTAC
CCCCGGCTATCGTGGCTACACCTACCGCCCCAGAAGACGAGCGACTACCCGACGCCGAACCACCACTGGAACCCGCC
GCCGCCGTCGCCGTCGCCAGCCCGTGCTGGCCCCGATTTCCGTGCGCAGGGTGGCTCGCGAAGGAGGCAGGACCCTG
GTGCTGCCAACAGCGCGCTACCACCCCAGCATCGTTTAAAAGCCGGTCTTTGTGGTTCTTGCAGATATGGCCCTCAC
CTGCCGCCTCCGTTTCCCGGTGCCGGGATTCCGAGGAAGAATGCACCGTAGGAGGGGCATGGCCGGCCACGGCCTGA
CGGGCGGCATGCGTCGTGCGCACCACCGGCGGCGGCGCGCGTCGCACCGTCGCATGCGCGGCGGTATCCTGCCCCTC
CTTATTCCACTGATCGCCGCGGCGATTGGCGCCGTGCCCGGAATTGCATCCGTGGCCTTGCAGGCGCAGAGACACTG
ATTAAAAACAAGTTGCATGTGGAAAAATCAAAATAAAAAGTCTGGAGTCTCACGCTCGCTTGGTCCTGTAACTATTT
TGTAGAATGGAAGACATCAACTTTGCGTCTCTGGCCCCGCGACACGGCTCGCGCCCGTTCATGGGAAACTGGCAAGA
TATCGGCACCAGCAATATGAGCGGTGGCGCCTTCAGCTGGGGCTCGCTGTGGAGCGGCATTAAAAATTTCGGTTCCA
CCATTAAGAACTATGGCAGCAAGGCCTGGAACAGCAGCACAGGCCAGATGCTGAGGGACAAGTTGAAAGAGCAAAAT
TTCCAACAAAAGGTGGTAGATGGCCTGGCCTCTGGCATTAGCGGGGTGGTGGACCTGGCCAACCAGGCAGTGCAAAA
TAAGATTAACAG
```
